# Compression-dependent microtubule reinforcement comprises a mechanostat which enables cells to navigate confined environments

**DOI:** 10.1101/2022.02.08.479516

**Authors:** Robert J. Ju, Alistair D. Falconer, Kevin M. Dean, Reto P. Fiolka, David P. Sester, Max Nobis, Paul Timpson, Alexis J. Lomakin, Gaudenz Danuser, Melanie D. White, Dietmar B. Oelz, Nikolas K. Haass, Samantha J. Stehbens

## Abstract

Cells migrating through complex 3D environments experience considerable physical challenges including tensile stress and compression. To move, cells need to resist these forces whilst also squeezing the large nucleus through confined spaces. This requires highly coordinated cortical contractility. Microtubules can both resist compressive forces and sequester key actomyosin regulators to ensure appropriate activation of contractile forces. Yet, how these two roles are integrated to achieve nuclear transmigration in 3D is largely unknown. Here, we demonstrate that compression triggers reinforcement of a dedicated microtubule structure at the rear of the nucleus by the mechanoresponsive recruitment of CLASPs (cytoplasmic linker-associated proteins) which dynamically strengthens and repairs the lattice. These reinforced microtubules form the mechanostat: an adaptive feedback mechanism that allows the cell to both withstand compressive force and spatiotemporally organise contractility signalling pathways. The microtubule mechanostat facilitates nuclear positioning and coordinates force production to enable the cell to pass through constrictions. Disruption of the mechanostat imbalances cortical contractility, stalling migration and ultimately resulting in catastrophic cell rupture. Our findings reveal a new role for microtubules as cellular sensors which detect and respond to compressive forces, enabling movement and ensuring survival in mechanically demanding environments.

**One Sentence Summary:** Mechanically tuned microtubules form a mechanostat to coordinate contractility and nuclear positioning in confined migration.

## Main Text

Cells live in confined environments where they must squeeze through narrow spaces, such as matrix pores, along tissue tracks and intercellular spaces, which presents a significant mechanical challenge. Very little is known about how cells sense and resist compression, despite it being a dominant mechanical input in cells in confined environments. As the largest and stiffest organelle, the cell nucleus is a major impediment to migrating through a confinement. To successfully transmigrate constrictions, cells must correctly position and protect the nucleus, whilst spatiotemporally coordinating actomyosin contractility to efficiently squeeze through the confinement. Microtubules are required for cell migration through mechanically confined 3D environments^1,2^, yet the reason for this remains unknown. Confinement restrains cell shape, crowding microtubule organization and inducing high curvature of polymers that physically damages the microtubule lattice^3^. Microtubules can adapt their biomechanical properties to resist compressive forces^4^ yet are pressure sensitive^5^. They exhibit self-repair behaviors^6^ by modifying the physical properties of microtubule polymers through the local incorporation of GTP-tubulin dimers at damage sites. Repaired polymers are more resistant to bending-induced breakage, resulting in longer lived, more stable microtubules. Mechanical forces are transmitted across the microtubule lattice, communicating the biophysical forces of the surroundings to intracellular structures physically linked to microtubules, including the nucleus^7, 8^. Importantly, microtubules sequester key upstream actomyosin regulatory factors^9^, to ensure appropriate timing of release and activation of contractile forces to facilitate cell shape changes and movement^10^. Thus, localised mechanoreponsive strengthening mechanisms are necessary to ensure that microtubules are reinforced in areas of high mechanical load. This facilitates both positioning and protection of organelles, as well as spatiotemporal regulation of the microtubule-contractility axis that controls migration^10, 11^. It remains unclear how microtubules are locally repaired and reinforced to drive cell motility in 3D environments.

### A microtubule cage in confined cells

To understand microtubule organization and function in 3D environments, we applied live-cell Light Sheet Fluorescence Microscopy (LSFM)^12^ and volumetrically imaged endogenously tagged microtubules (α-tubulin-meGFP; CRISPR/Cas9 edited) in a highly migratory cell line, 1205Lu melanoma cells, embedded deep in reticulated collagen hydrogels (Fig. 1a). We observed that in comparison to 2D (Extended Data Fig. 1a), microtubules in these conditions were organised in a highly curved and dynamic cage-like structure enclosing the nucleus and lining the cell cortex (Fig. 1b, Extended Data Fig. 1b, c; Supplementary Video 1 and 2). As the cell routed between reticulated collagen constrictions, the cage dynamically reorganised, the nucleus constantly repositioned and microtubules accumulated at points of cellular constriction (Fig. 1b). Microtubules that exhibit high curvature are prone to lattice fractures and depolymerization^3^. Thus, we hypothesised that the maintenance of this highly curved cage-like structure may require mechanical reinforcement through dynamic repair mechanisms.

**Fig.1.**
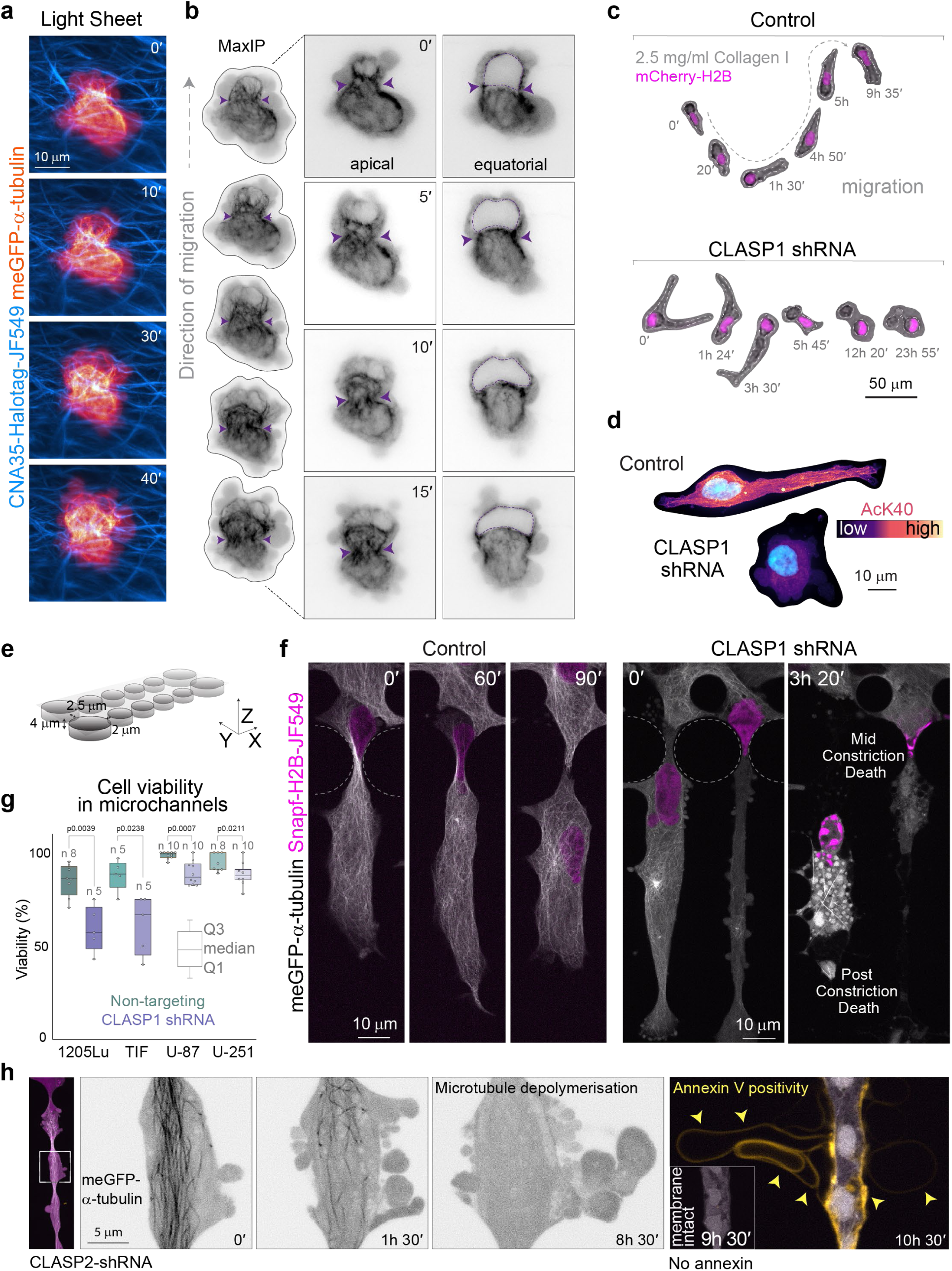
ACLASP1-dependent dynamic microtubule cage is required for cell migration in complex 3D environments. (**a**) 1205Lu cells endogenously expressing meGFP-α-tubulin migrating through a reticulated 3D collagen I hydrogel. Light sheet fluorescence microscopy (LSFM) volumetric imaging reveals microtubule structural dynamics as cells squeeze between matrix fibres. (**b**) Microtubule organisation (meGFP-α-tubulin, contrast inverted) during constriction (purple arrowheads). Apical and equatorial Z sections of cell in **a**. Purple dotted line defines nucleus. (**c**) CLASP1 depletion causes cell death during 3D migration. Representative time-lapse of control (non-targeting) and CLASP1-depleted nuclear labelled (mCherry-H2B) 1205Lu cells navigating 3D collagen I hydrogels. (**d**) Immunofluorescence of acetylated alpha-tubulin (AcK40) in control (non-targeting) and CLASP1-depleted cells in collagen I hydrogels. (**e**) PDMS Constriction microchannel design. Cylinders represent PDMS pillars forming constrictions at defined spacings. (**f**) Spinning disc confocal microscopy time-lapse of 1205Lu cells with fluorescently labelled microtubules (meGFP-α-tubulin) and nuclei (Snapf-H2B-JF549) in constriction microchannels. Constriction pillars marked by grey dotted line. CLASP1 depletion results in cell rupture and death. (**g**) CLASP1 depletion results in significantly more cell death in microchannels over 12 hours (1205Lu melanoma, TIF telomerase-immortalised fibroblasts, U-87 and U-251 glioblastoma). (**h**) CLASP2-depleted 1205Lu cell endogenously expressing meGFP-α- tubulin (grey) and nuclear marker, 3X NLS mScarlet-I (magenta) incubated with Annexin V-647 (yellow). Magnified region from left panel (contrast inverted): microtubule depolymerisation coincides with excessive membrane blebbing (1h 30’-8h 30’) and precedes annexin positivity (9h-30’-10h 30’).

### CLASP-mediated reinforcement of the microtubule cage allows cells to withstand confinement

To investigate the stability of the microtubule cage, we examined mechanisms that repair damaged microtubules in response to mechanical insult. Cytoplasmic linker-associated proteins (CLASPs) are a family of plus-end tracking proteins (+TIPs) that act as microtubule rescue or anti-catastrophe factors, spatio-temporally regulating microtubule growth^13^. +TIPs such as CLASP, support the continual growth of microtubules, which enhances acetylation of the intraluminal Lysine 40 residue of α-tubulin (AcK40) - stabilizing microtubules^14^ and making polymers more flexible in response to mechanical force^15^. CLASPs also assist repair of mechanical damage to polymers by locally exchanging tubulin dimers at fracture sites^16–18^. Strikingly, when CLASPs were depleted, cells navigated tight spaces in 3D collagen more slowly and eventually ruptured and died (Fig. 1c, Extended Data Fig. 2a–f). Furthermore, microtubule acetylation was lost in CLASP-depleted cells in 3D collagen (Fig. 1d, Extended Data Fig. 3a) consistent with reports in 2D^19^ (Extended Data Fig. 3b, 2a), with no detectable change in total tubulin levels (Extended Data Fig. 3c). To control the spatial and mechanical constraints experienced by migrating cells, we microfabricated confinement channels with constrictions consisting of cylindrical pillars (2-2.5 µm x 4 µm) representing pore sizes cells navigate through in 3D collagen gels (Fig. 1e, Extended Data Fig. 4a)^20, 21^. Using 3D Structured Illumination Microscopy (3D-SIM) to volumetrically image cells in microchannels we confirmed that cells similarly form a microtubule cage (Supplementary Video 3; z-scroll). As in 3D collagen, CLASP-depleted cells rupture in confinement (Fig. 1f, Extended Data Fig. 4b–c, Supplementary Video 4). This behaviour was consistent across melanoma (1205Lu), glioblastoma (U-87, U-251) and fibroblast cell lines (TIF) (Fig. 1g, Extended Data Fig. 4b). Again, CLASP-depletion caused microtubule depolymerisation specifically when cells experienced confinement preceding annexin V positivity (Fig. 1h) suggesting microtubules are required to resist cell death induced by mechanical forces. These data demonstrate that in the absence of CLASPs, microtubules are unable to resist mechanical compression and depolymerise, resulting in cell rupture and death.

**Fig.2.**
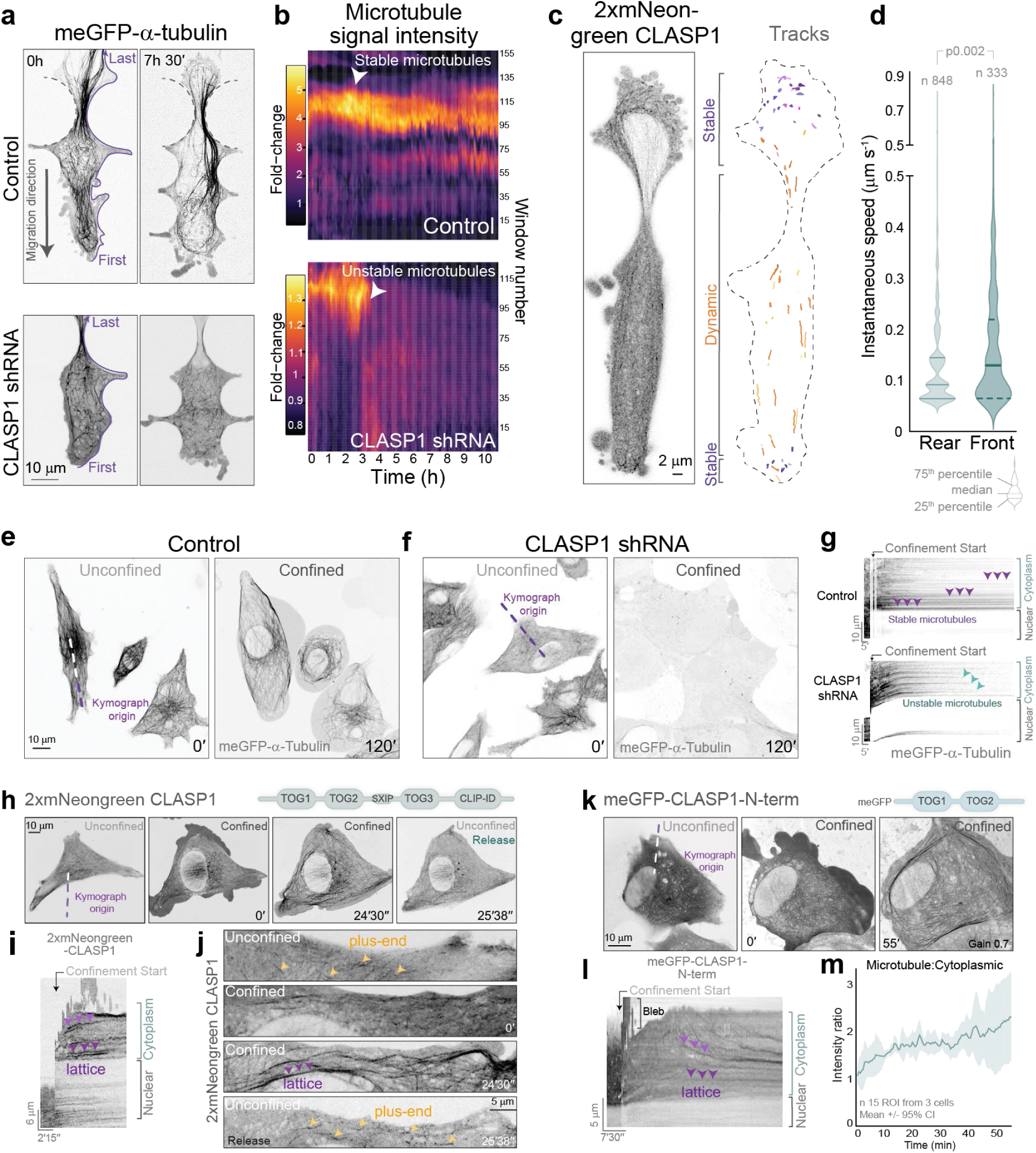
CLASPs spatiotemporally regulate microtubule mechanical stability in confined environments. (**a**) CLASP1-depleted microtubules destabilise upon mechanical constriction in microchannels. Representative time-lapse of control (non-targeting) and CLASP1-depleted 1205Lu cells endogenously expressing meGFP-α-tubulin. Quantitation windows dynamically wrap with cellular edge over 11 hours (purple line shows window placement). (**b**) Kymographs of quantitation windows in **a** demonstrate abrupt microtubule loss in CLASP1-depleted cells. (**c**) 1205Lu cell expressing 2xmNeonGreen-CLASP1 migrating through a microchannel constriction. CLASP1 dynamics over 45 seconds reveal distinct pools of stable (purple) and dynamic (orange) microtubule association. (**d**) CLASP1 dynamics (instantaneous speed) in rear and front cellular compartments during constriction. (**e**) Representative time-lapse of control (non-targeting) and (**f**) CLASP1-depleted 1205Lu cells endogenously expressing meGFP-α-tubulin, before and during axial confinement. (**g**) Kymographs of meGFP-tubulin demonstrate a loss of microtubules in CLASP1-depleted cells. (**h**) 2xmNeonGreen-CLASP1 switches localization between microtubule plus-ends and the lattice upon dynamic axial confinement. (**i**) CLASP Kymograph of region in **h**. (**j**) Higher magnification of CLASP1 mechano-dynamics in axial confinement. (**k**) Representative time-lapse of meGFP-CLASP1-N-term (TOG1 and TOG2) compression induced microtubule loading. meGFP-CLASP1-N-term is cytoplasmic in unconfined conditions. (**l**) CLASP Kymograph of region in **k**. (**l**) Average meGFP-CLASP1-N-term microtubule association dynamics. eGFP microtubule:cytoplasmic fluorescence intensity ratio profile, measured as a function of time.

### Mechanoresponsive CLASP localization ensures spatiotemporal microtubule stability

Microtubule repair factors such as CLASPs and microtubule acetylation, act as proxy signals to label sites of microtubule lattice damage and repair^18, 22^. We hypothesised that these repair factors would be recruited to cellular regions experiencing mechanical tensile or compressive stress to locally repair and reinforce the microtubule lattice. We confirmed that CLASPs were required for microtubule stability in microchannel confinement (Fig. 2a, b). Next, we examined the localization of CLASP1 (2xmNeonGreen-CLASP1) as cells passed through constrictions. After the front of the cell enters a constriction, the nucleus becomes occluded and must deform to transmigrate through the confined region (Extended Data Fig. 4d). We found that once nuclear occlusion occurred, CLASPs were spatially regulated localising to distinct pools: 1) a near static (stable) rear pool in the cellular compartment behind the nucleus, and at the leading edge of the cell, and 2) a dynamic pool associating with growing plus-ends in the compartment in front of the nucleus (Fig. 2c, d, Extended Data Fig. 5a–c). As the rear cytoplasmic compartment shrinks and compresses microtubules against the occluded nucleus, this likely induces lattice fractures and the stable recruitment of CLASPs.

To confirm that compressive forces recruit CLASPs to microtubules we used an acute axial confinement approach^23^. CLASP1 was required to stabilise microtubules (Fig. 2e–g, Supplementary Video 5) and maintain membrane integrity (Extended Data Fig. 6), verifying our microchannel results. Acute compressive confinement of 2xmNeonGreen-CLASP1 expressing cells induced a shift in CLASP1 localisation from microtubule plus-ends to along the lattice, signifying that CLASP1 localisation is responsive to compressive force (Fig. 2, h-j, Supplementary Video 6; 3 microns axial confinement). This mechanoresponse was dependent on the direct N-terminal TOG1 and TOG2 interaction with microtubules (Fig. 2, k-m, Extended Data Fig. 7). This suggests that nuclear occlusion causes compressive forces on the microtubules in the rear compartment leading to the recruitment of CLASPs.

We hypothesised that the recruitment of CLASPs stabilises the microtubules in the rear compartment. Therefore, we examined the post-translational tubulin modifications AcK40 and Tyr that identify distinct populations of stable and dynamic microtubules, respectively^14, 24^. A population of stable AcK40-positive microtubules was enriched in the rear compartment forming a cushion-like structure behind the occluded nucleus (Fig. 3a; early, 3b; control, Extended Data Fig. 8a, b, Supplementary Video 3). As cells progressed through constrictions, the rear microtubule cushion disappeared and the AcK40 localised to parallel microtubule bundles enveloping the nucleus (Fig. 3a; mid) – a similar location to where Arp2/3-dependent actin nucleation generates lateral pressure on the nucleus to deform it^25^. Once the nucleus progressed through a constriction, the spatial restriction of the AcK40 was lost (Fig. 3a; post/pre). During nuclear occlusion of CLASP-depleted cells, the microtubule cushion failed to form, with no accumulation of AcK40 in the rear compartment (Fig. 3c). The absence of the cushion was concurrent with uncaged and mispositioned nuclei (Extended Data Fig. 8c). This suggested the spatiotemporal association of lattice repair factors act to stabilise the microtubule cushion and position the nucleus for transmigration.

**Fig.3.**
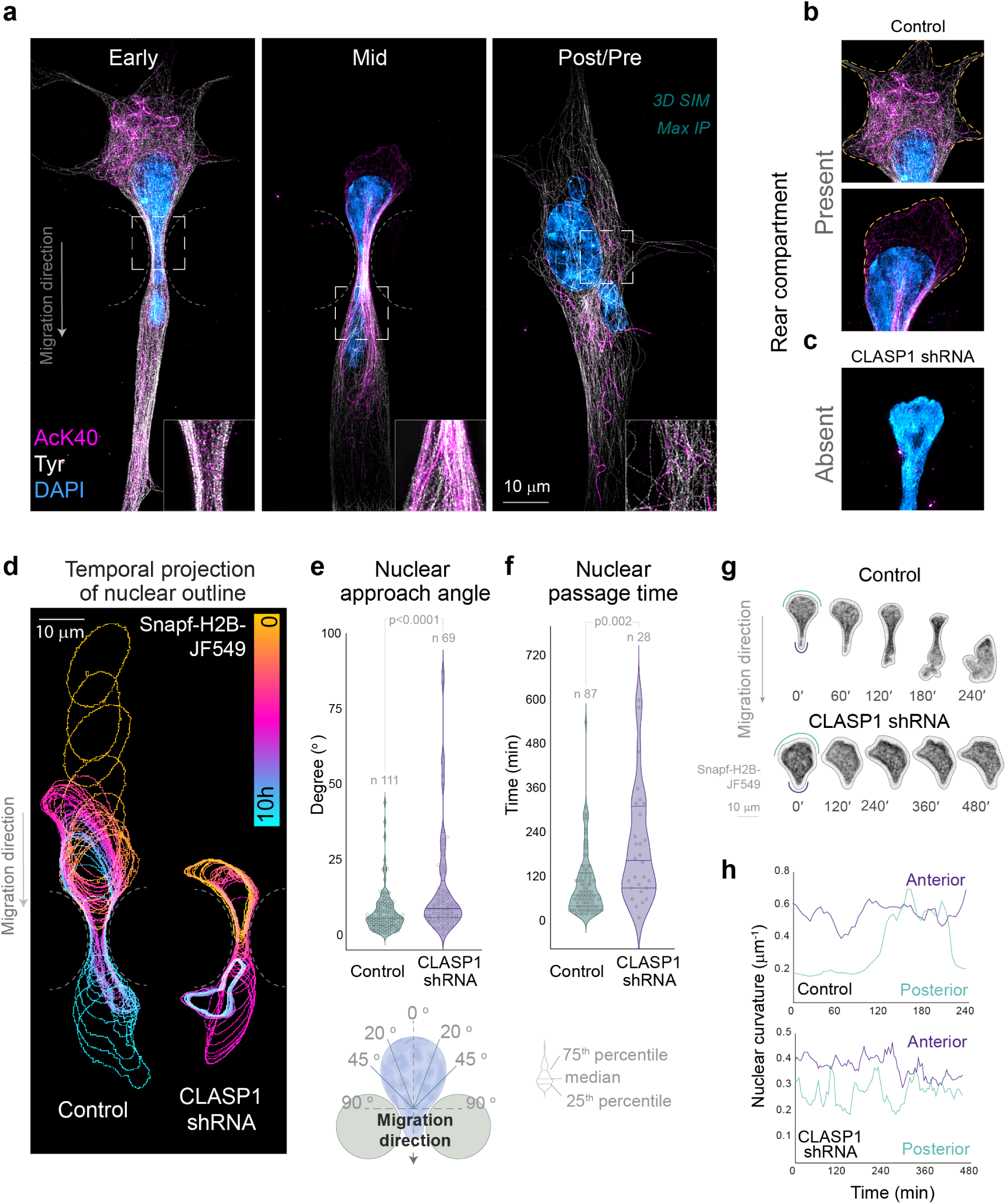
CLASP1-reinforced microtubules are required to position and protect the nucleus in confined cells. (**a**) 3D-Structured Illumination microscopy of control (non-targeting) 1205Lu cells early, mid and post/pre-constriction. Acetylated (AcK40) and tyrosinated α-tubulin (Tyr) immunolabelled 1205Lu cells, nuclei labelled with DAPI. (b) Magnified regions from control and (**c**) CLASP1-depleted cells undergoing confinement. CLASP1-depleted cells lack microtubules in the rear compartment. (**d**) Temporal colour-coded projection of nuclear outlines of control (non-targeting) and CLASP1-depleted cells undergoing constriction in microchannels. (**e**) Quantitation and schematic of nuclear approach angle in control and CLASP1-depleted cells. (**f**) Analysis of nuclear passage time in control and CLASP1-depleted cells. (**g**) Representative time-lapse images of Snapf-H2B-JF549I labelled nuclei, control, and CLASP1-depleted cells. (**h**) Graph of nuclear curvature of anterior and posterior surfaces in cells undergoing constriction. Anterior nuclear curvature remains stable, whereas posterior nuclear curvature increases only as nucleus progresses through constriction. CLASP1 depletion alters nuclear curvature dynamics.

### CLASP-reinforced microtubules are required for nuclear positioning during confinement

As cells entered constrictions, they positioned their nucleus (Snapf-H2B-JF549) with the long axis aligned with the axis of migration (Fig. 3d, e, Extended Data Fig. 9a). However, CLASP-depleted cells failed to reorient their nuclei (Fig. 3, d and e) and spent significantly longer in highly constricted states of non-productive migration (Fig. 3f, Extended Data Fig. 9a, b) with nuclear deformation (Fig. 3g, h, Extended Data Fig. 9c). CLASP-depleted cells showed an increase in nuclear rupture events compared to control cells (Extended Data Fig. 9d), indicating that the loss of the microtubule cushion also results in nuclear damage. Thus, CLASP-reinforced microtubules are required for nuclear protection and positioning to facilitate efficient nuclear transmigration.

### CLASP-stabilised microtubules are required to balance cortical contractility and hydrostatic pressure

CLASP-depleted cells undergoing constricted migration display persistent, abnormal cortical blebbing (Fig. 1h, Supplementary Video 4). Blebs are membrane protrusions that form when the membrane detaches from the underlying cortical actin. Influx of cytosolic content enlarges the membrane protrusion, which can retract once the pressure is re-equilibrated and balanced by repair of the cortex^26, 27^. In constricted control cells with fluorescently labelled membranes (LCK-mScarlet-I) and F-actin (Lifeact-eGFP), we observed occasional small blebs at the rear membrane (Fig. 4a, b) potentially indicating higher cortical contractility and hydrostatic pressure in the rear compartment than in the front. This polarised rear blebbing quickly resolved once cells progressed through constrictions (Supplementary Video 7). By comparison, constricted CLASP-depleted cells exhibited significantly larger, more persistent blebs with faster protrusion velocities, in both the front and rear of cells (Fig. 4a–c, Extended Data Fig. 10a–d, Supplementary Video 7). This unpolarised blebbing behaviour persisted for prolonged periods during which cells could not progress through constrictions (Supplementary Video 7). Whilst the blebs in CLASP-depleted cells protruded with a greater velocity than control cells, they resolved at the same retraction velocity (Extended Data Fig. 10d) indicating that cortical repair mechanisms were intact. These data suggest that mechanically reinforced microtubules are required to locally harmonise cortical contractility and hydrostatic pressure.

**Fig.4.**
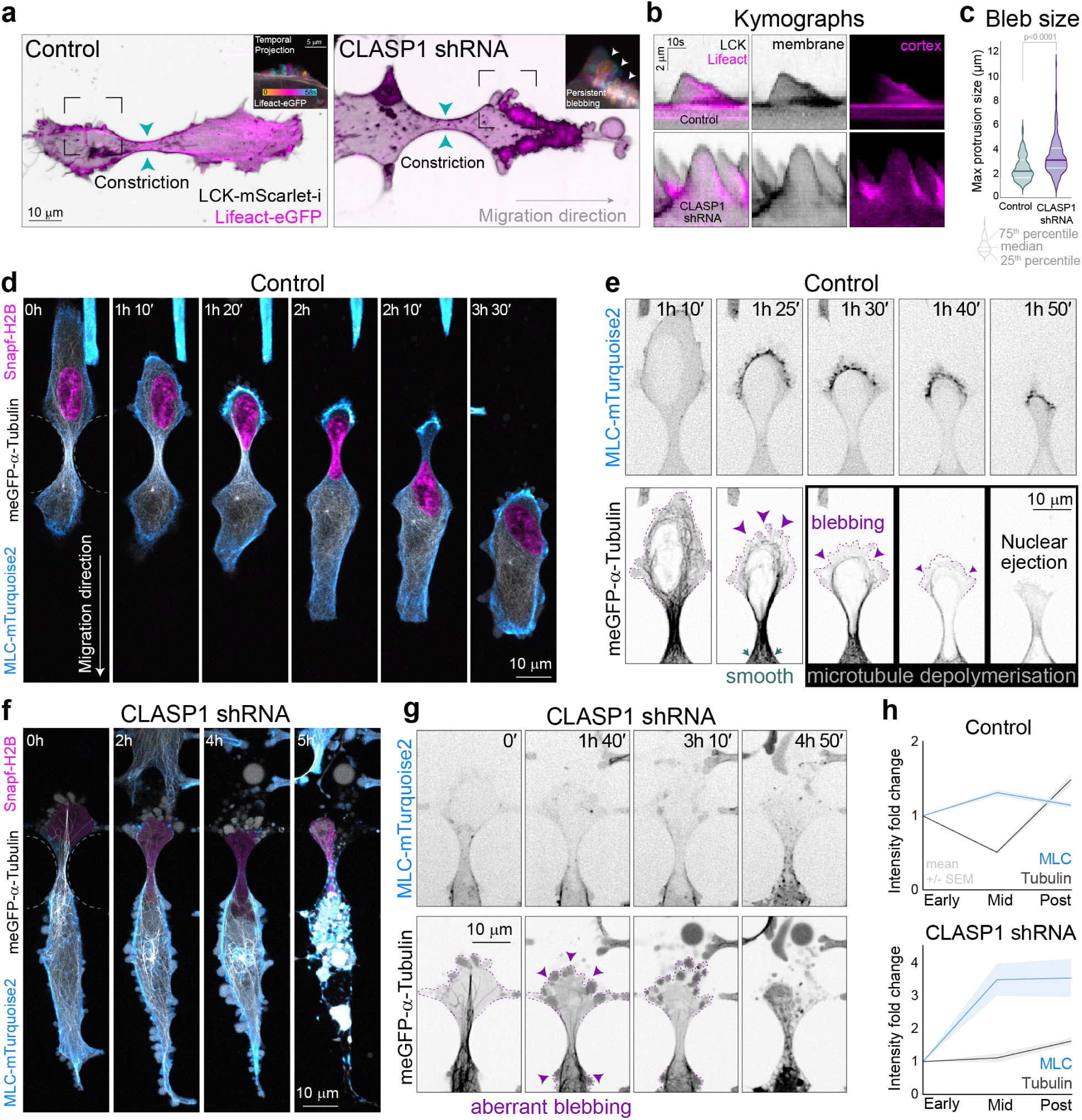
CLASP1-reinforced microtubules are required to coordinate contractility in confined cells. (**a**) Time-lapse images of control (non-targeting) and CLASP1-depleted cells co-expressing membrane (LCK-mScarlet-I) and cortical (eGFP-Lifeact) markers undergoing nuclear constriction in microchannels. Insets are temporal colour-coded projections (1 frame/sec for 1 min) of actin bleb dynamics at the cell surface. CLASP1-depletion results in persistent pericellular blebbing during cellular constriction (temporal projection inset). (**b**) Kymographs of membrane (black) and cortical (magenta) dynamics in control and CLASP1-depleted cells. (**c**) CLASP1-depletion increases bleb size. Time-lapse images of control (**d-e**) and CLASP1-depleted (**f-g**) 1205Lu cells with fluorescently labelled microtubules (meGFP-α-tubulin), nucleus (SnapTag-H2B-JF549) and myosin light chain (MYL9, MLC-mTurquoise2) undergoing constricted migration. Contrast inverted images (**e**) demonstrate myosin concentration on the rear membrane, proximal to the microtubule filled compartment (meGFP-α-Tubulin; microtubule ‘cushion’) which correlates with an increase in membrane blebbing (purple dotted line and arrowheads) as microtubules disassemble. (**g**) Myosin does not concentrate to the rear compartment of CLASP1-depleted cells. (**h**) Quantitation of myosin and microtubule fluorescence intensities over time in the rear cellular compartment during nuclear translocation in control and CLASP1-depleted cells.

### The microtubule cushion is required for spatiotemporal control of contractility for nuclear transmigration

We then sought to understand how CLASP-dependent microtubule reinforcement might locally control actomyosin contractility. First, we investigated the spatiotemporal relationship between microtubules and myosin during nuclear transmigration. As control cells labelled for microtubules, myosin light chain and nuclei (eGFP-α-tubulin, MLC-mTurquoise2 and Snaptag-H2B respectively), entered constrictions and the nucleus became occluded, myosin accumulated proximal to the rear plasma membrane (Fig. 4d, Supplementary Video 8) consistent with previous studies^23, 25, 28–30^. The myosin signal increased in intensity as the nuclear constriction progressed and the rear compartment membrane contracted maximally. The microtubule cushion formed by highly curved microtubules, filled and maintained the cytoplasmic compartment between the nucleus and rear membrane (Fig. 4e, Supplementary Video 8). Although the microtubules in the front cellular compartment maintained a polymer structure, the rear microtubule cushion disassembled immediately prior to nuclear propulsion through the constriction (Fig. 4e; purple arrowheads, Fig. 4h, Supplementary Video 8). Consistent with our earlier observation (Fig. 3b, Extended Data Fig. 8c), this rear pool of mechanically reinforced microtubules was noticeably absent in CLASP1-depleted cells (Fig. 4f–h, Supplementary Video 8). We propose that the cushion of CLASP-reinforced microtubules forms in response to increasing hydrostatic pressure arising from contraction of the rear membrane. The cushion allows the hydrostatic pressure behind the nucleus to reach a critical point to overcome the resistance to nuclear translocation^29, 30^.

Microtubules regulate the spatial and temporal activation of Rho-dependent actomyosin contractility of cells^10^, a phenomenon recently named the ‘microtubule-contractility axis’^11^. The RhoA activating protein GEF-H1 (ARHGEF2), is a microtubule-bound GEF which locally integrates microtubule and actin dynamics^31, 32^. GEF-H1 is held in a catalytically inactive state on microtubule polymers, and is activated upon release from depolymerising microtubules^33^. To investigate if mechanically reinforced microtubules control myosin localisation through sequestration and release of GEF-H1, we imaged cells expressing a GEF-H1 activity biosensor (GEF-H1 FLARE212)^32^ (Fig. 5a, Supplementary Video 9). Following nuclear occlusion in constrictions, GEF-H1 activity rapidly increased in the rear compartment due to release of inactive GEF-H1 from the depolymerising microtubule cushion (Fig. 5a; high FRET, yellow, Extended Data Fig. 10e, f). The nucleus then rapidly passed through the constriction. Consistent with our observations of myosin localisation, CLASP-depleted cells which lack the microtubule cushion were unable to concentrate GEF-H1 activity to the rear of cells (Fig. 5a, b; CLASP1shRNA). Furthermore, cells undergoing nuclear transmigration revealed a burst of Rho (eGFP-RhoA)^34^ behind the nucleus (Fig. 5c, d, Supplementary Video 10) correlating with the timing of microtubule depolymerization (Fig. 4e) and GEF release (Fig. 5a, b) in the rear compartment immediately prior to nuclear translocation. This coordinated Rho accumulation was also absent in CLASP-depleted cells (Fig. 5c, d; CLASP1shRNA). Instead, Rho accumulation fluctuated at various locations across the cell membrane in both the front and rear compartments over many hours (>8 h). These aberrant Rho dynamics were coincident with non-productive migration where cells would oscillate back and forth in a constricted state (Supplementary Video 10). Thus, mechanical reinforcement of the microtubule cushion by CLASPs enables spatiotemporal control of the microtubule- contractility axis, allowing cells to efficiently navigate through confinement.

**Fig.5.**
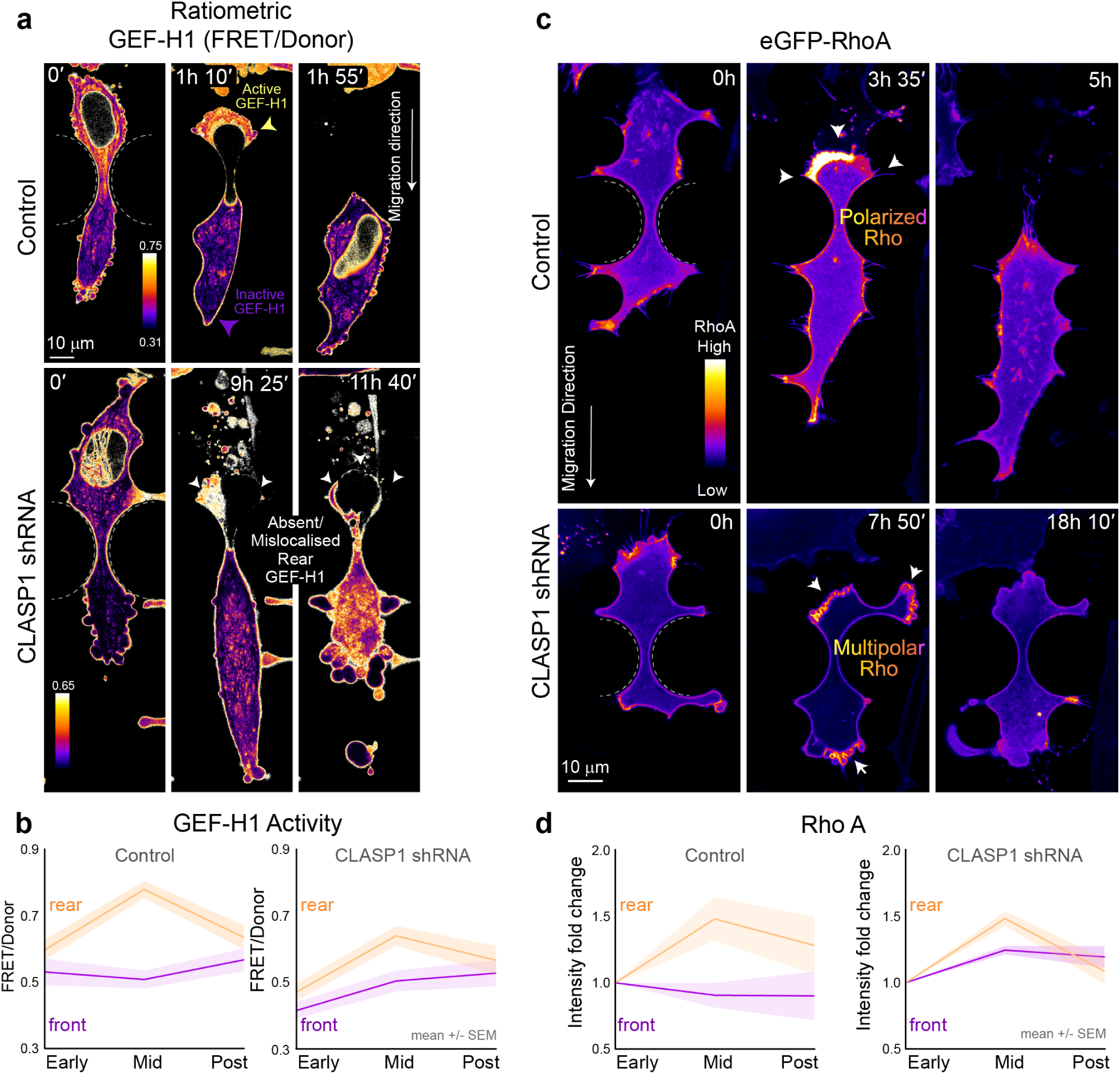
Controlled microtubule depolymerisation is required to spatially restrict contractility signalling. (**a**) Time lapse images of GEF-H1 FLARE212 localization in control (non-targeting) and CLASP1-depleted cells undergoing constriction. GEF-H1 activation (Control; yellow, high FRET/donor ratio) arising from microtubule depolymerisation is polarised posterior to the nucleus during nuclear translocation (1 h 10 min). The polarisation of active GEF-H1 equilibrates after cells translocate through constrictions (1 h 55 min), shown by inactive (purple, low FRET/donor ratio) GEF-H1 association with microtubules. GEF-H1 activation is aberrantly localised in CLASP1-depleted cells. Cells subsequently spend longer in constricted states (0 – 9 h 25 min), resulting in cell rupture (11 h 40 min). (**b**) Dynamics of GEF-H1 FRET activity as a function of nuclear constriction phases-in front (purple) and rear (orange) compartments of control and CLASP1-depleted cells. (**c**) Rho-A accumulates posteriorly prior to a Rho-burst (3 h 35 min; arrowheads) which completes nuclear ejection through translocation (5 h). In CLASP1-depleted cells Rho-A no longer concentrates at the rear and occurs both posteriorly and anteriorly (arrowheads) with erratic localization at large blebs. (**d**) Dynamics of Rho-A intensity as a function of nuclear constriction phases-in front (purple) and rear (orange) compartments of control and CLASP1-depleted cells.

### Theoretical model of a microtubule mechanostat in confined cell migration

We formulated a mathematical model for single-cell transmigration between two micro-pillars to understand how microtubules spatiotemporally control contractility during nuclear transmigration (Fig. 6a). In this model, both the cell cortex and the nuclear envelope are represented by closed 2D contour lines. When the nucleus is occluded it establishes front and rear cytoplasmic compartments. The mechanical component of the model relies on the balance of a constant migratory force, friction, viscous and elastic forces of nuclear centring, cortical tension, cytoplasmic hydrostatic pressure as well as nuclear envelope tension and nuclear pressure (Fig. 6b). The mechanical model is complemented by rearward cortical myosin flow, leading to an increase in contractility and rear-to-front flux of cytoplasm in response to the increase in rear pressure. We also involve a dense network of mechanically reinforced microtubules in the rear cytoplasmic compartment acting as the ‘cushion’ – preventing the rear compartment from reaching a zero volume (blue area in Fig. 6a). When the rear pressure exceeds the cushion’s resistance to compression, we model the disassembly of microtubules (Extended Data Fig. 11a). This results in a burst of Rho activity which increases cortical tension in the rear compartment (Fig. 6c).

**Fig.6.**
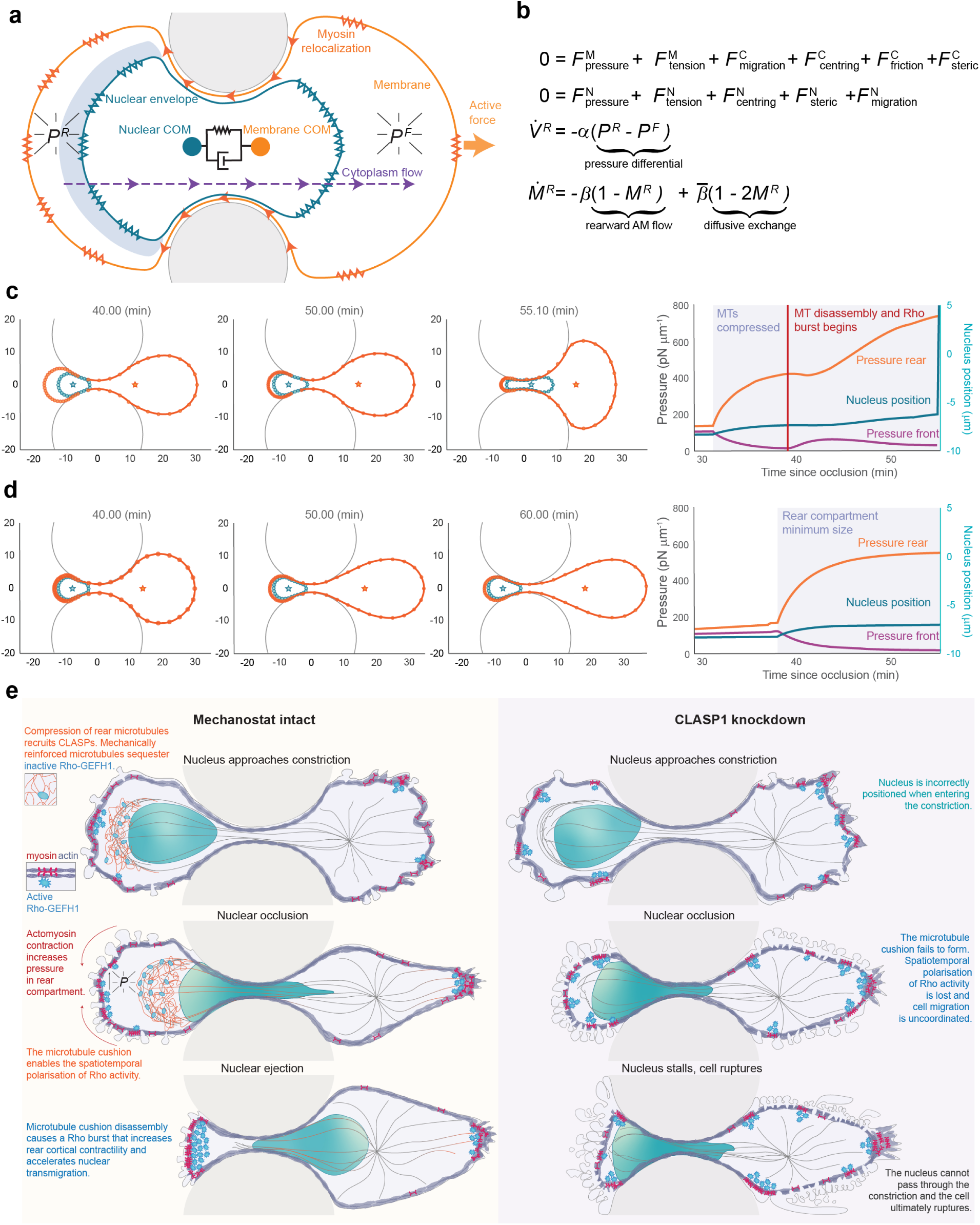
Theoretical model of mechanostat function in confined cell migration. (**a**) A system of force balance equations governs the shape and motion of cell and nucleus. Tension in the actomyosin cortex and the nuclear envelope increase the rear pressure PR and front pressure PF in the associated compartments. A Kelvin-Voigt element (elastic spring and viscous dash-pot in parallel) between the centres of mass of the cell and the nucleus, models nuclear positioning. (**b**) Rate equations describe myosin aggregation in the rear due to the rearward actomyosin flux and diffusive exchange, as well as forward flux of cytoplasm during occlusion. (**c**) The simulated progression of a cell through the constriction: The pressure graph shows an initial pressure differential upon first entering the occlusion which increases once the volume of the rear cytosolic compartment is stabilised by the microtubule cushion (shaded area). Gradual up-regulation of rear cortical tension in response to a burst of Rho activity following microtubule disassociation accelerates the build-up of pressure and of the velocity of the nucleus and leads to the passage of the nucleus past the pillars within minutes; (MT microtubule). (**d**) In the absence of a rear microtubule cushion and of myosin up-regulation, the rear cortex physically contacts the nuclear envelope during the attempted transmigration. The nucleus does not progress through the constriction within the simulated period of time. (**e**) Schematic summarizing the role of the microtubule mechanostat in confined migration.

Our simulations reveal that transmigration is initially prevented by the elastic resistance of the nucleus. Once occlusion occurs, myosin accumulates in the rear cortex, triggering a difference in cortical tension and a subsequent imbalance in cytoplasmic hydrostatic pressure between the front and the rear compartments (Fig. 6c). During this process the microtubule cushion is compressed-preventing close physical contact between the cortex and the nucleus-allowing for pressure to build in the rear. The microtubule cushion then disassembles, releasing a burst of Rho activity (Fig. 6c graph) that accelerates the build-up of tension in the rear cortex and increases the velocity of nuclear transmigration (Extended Data Fig. 11b, Supplementary Video 11). In simulations lacking the rear microtubule cushion and subsequent myosin up-regulation in response to microtubule disassembly, the nucleus fails to pass the constriction (Fig. 6d, Extended Data Fig. 11c, Supplementary Video 11). If sufficient pulling forces remain the nucleus passes at a much later time point – spending longer in a constricted state (Supplementary Video 11). Our theoretical modelling supports the concept of an adaptive feedback mechanism whereby microtubules are reinforced in response to compression, allowing cells to withstand force and spatiotemporally organise contractility signalling pathways. Analogous to the influence of mechanical loading on bone structure^35^, we define this feedback mechanism as a microtubule mechanostat (Fig. 6e).

## Discussion

Cells *in vivo* must migrate through crowded 3D environments where they become confined and are subject to compressive forces. We have established that microtubules are reinforced by a CLASP-dependent feedback mechanism in response to compression. These mechanically tuned microtubules act as a mechanostat that is dynamically repaired and required to coordinate cortical contractility and nuclear positioning for migration.

Nuclear deformation and occlusion within tight pores divides the cell into front and rear compartments. It has been proposed that nuclear occlusion can generate a gradient of intracellular pressure; low at the rear and high at the front^36^. However, more recent chemico-mechanical models suggest that cells overcome the resistance of nuclear constriction by a combination of actin contraction and cytosolic pressure from the rear^37^. The importance of a rear cytosolic hydrostatic pressure gradient has been shown to be integral in compressing the nucleus to promote constricted nuclear transmigration ^25, 28, 29, 37, 38^. While we know that this cytosolic back pressure requires locally focused actomyosin contractility, how this force-producing machinery is actively controlled at the rear within complex geometries is unclear. Our data demonstrate the local reinforcement of microtubules by CLASPs comprises a mechanostat that enables cells to 1) correctly position the nucleus and 2) spatiotemporally coordinate the actomyosin contractility that generates the cytosolic hydrostatic pressure required to overcome the nuclear occlusion. We propose that upon reaching sufficient cytosolic hydrostatic pressure in the rear cellular compartment, the microtubule cushion begins to disassemble and releases GEF-H1. This activates a final burst of RhoA-mediated contractility at the rear membrane which increases nuclear velocity, enabling the nucleus to escape the constriction.

The exact mechanism that causes the microtubule disassembly remains to be determined. Increasing cytosolic hydrostatic pressure may directly induce microtubule depolymerization^5^, potentially by inducing microtubule lattice packing defects that alter CLASP’s association with the polymer. Another formal possibility is that cytosolic hydrostatic pressure indirectly affects microtubule stability, potentially downstream of mechanosensitive ion channels^39^. Additionally, nuclear occlusion may limit the diffusion of GTP, tubulin or other microtubule-associated proteins required for microtubule self-repair to the rear compartment of cells. In the absence of reinforcement by CLASPs, microtubules may be less able to withstand the effects of increasing cytosolic hydrostatic pressure, causing inappropriate disassembly, release of GEF-H1 and uncoordinated actomyosin contractility. The inability to repair lattice damage and resist mechanical stress, likely drives a switch from a tightly regulated microtubule-contractility axis to an induction of global contractility. This results in a loss of front-rear polarity and an imbalance of the forces required for nuclear translocation.

Our findings demonstrate that the mechanostat is crucial for maintaining cellular mechanical integrity in confinement. We demonstrate a requirement for mechanoresponsive reinforcement of the microtubule lattice in response to compressive forces during constriction. Failure to efficiently repair and reinforce spatially distinct pools of microtubules during mechanical challenges promotes catastrophic cell rupture and death in confined 3D environments. Beyond its relevance in confined migration, the microtubule mechanostat may act in diverse settings where cells need to maintain their shape under compressive load. For example, in cardiac and skeletal muscle cells microtubules act as shock absorbers, forming load-bearing spring elements to control cell contractility and muscle strength^40, 41^. In articular cartilage, microtubules maintain chondrocyte cell volume and recovery in response to compression^42^. Thus, it is plausible that the microtubule mechanostat also reinforces polymers in these processes. Furthermore, as rational manipulation of the microtubule contractility axis results in T-cells that move through tissues faster^11^, tuning the mechanostat could improve treatments for cell-based immune therapies. Exploring the mechanostat’s role in different biological contexts will reveal how cells sense and respond to compressive forces during development and disease and inform therapeutic microtubule targeting strategies.

## Methods

### Cell Culture

1205Lu human melanoma cells^43^, Telomerase-immortalised fibroblasts (TIFs)^44^, glioblastoma astrocytoma cell lines U-251 MG (Merck, SKU09063001-1VL) and U-87 MG (Merck, SKU 89081402-1VL), HEK293FT cells (ThermoFisher Scientific, R70007) were all maintained in high glucose DMEM (Gibco, 11965092), 10% foetal bovine serum (Gibco, 10100147), 0.1 mM MEM non-essential amino acids (Gibco, 11140050) and 100 i.u. mL^−1^ penicillin, 100 μg mL^−1^ streptomycin (Gibco, 15070063). 293FT cells were maintained in 0.5 mg mL^-1^ G418 (Gibco, 10131-035). Cells were routinely subjected to PCR mycoplasma testing^45^ and confirmed negative.

3D Atelo Bovine collagen I (Advanced Biomatrix, #5133, FibriCol) hydrogels were prepared by diluting collagen I stock to a final concentration of 2.5 mg/mL with 10% v/v 10x MEM-α (BE12-684F, Lonza), 10% v/v foetal bovine serum, 2 mM L-glutamine (Gibco, 25030) and sterile water. Hydrogel solutions were titrated to ∼pH 7 using sodium bicarbonate (Gibco, 25080094) with pH indicator strips (Merck, 109535). Collagen I solution preparation was performed on ice with all reagents pre-chilled to 4°C, polymerisation was induced by incubation at 37°C, 5% CO_2_ in a tissue culture incubator for 30-60 minutes after cell embedding, before imaging. Recombinant CNA35-Halotag collagen I binding peptide was purified and used as previously described^46^, endotoxin levels were reduced using spin columns (Thermo Fisher Scientific, 88274,). DQ™-collagen was used as per manufacturer’s instructions (Invitrogen, D12060).

### DNA constructs and lentiviral vectors

All plasmids will be made available on Addgene and upon reasonable request. To generate stable fluorescent protein expressing 1205Lu cell lines, proteins of interest were subcloned into entry vectors and recombined into lentiviral destination vectors-except for the eGFP-RhoA expressing cells. Stable 1205Lu-RhoA cells were generated as described previously^47^. The RhoA construct^34^, was cloned into the pPB-CAG.EBNXN vector^48^ by NdeI/SalI excision from its pcDNA3.1 backbone. pPB-CAG.EBNXN-RhoA-FRET was then co-transfected with pCMV-hyPBase^49^ into 1205Lu cells. Changes in RhoA localisation during constricted migration were calculated using eGFP fluorescence alone.

Entry vectors were created as follows. pENTR-SNAPf-H2B was created by restriction digest of pENTR1a-GFP-N2 (FR1) (Addgene, #19364 Eric Campeau and Paul Kaufman) and pSNAPf-H2B control plasmid (Addgene #101124, New England Biolabs and Ana Egana) using KpnI and NotI, and ligated together using T4 DNA ligase (New England Biolabs, M0202). The following constructs were created using HiFi assembly (New England Biolabs, E2521L). Compatible fragments were combined as per manufacturer’s instructions.

pENTR-3xNLS-mScarlet-I-nucleoplasmin-NLS was created using HiFi assembly (New England Biolabs). The 3xNLS-mScarlet-I insert was synthesised as a gblock (Genewiz) and PCR amplified (New England Biolabs, M0515, Q5 Hot start) with primers FW 5’- CCACCGGCCGGTCGCATGCCAAAAAAGAAGAGAAAGGTAGACC -3’ and RV 5’- TCGAGTGCGGCCGCTTTATTTCTTTTTCTTAGCTTGACCAGCTTTCTT -3’ to create overhanging microhomologies. A microhomology compatible vector was created by PCR amplification of pENTR1a-eGFP-N2 (FR1), (Addgene, #19364 Eric Campeau and Paul Kaufman) using the primers ENTR-FW 5’-AGCGGCCGCACTCGAGA-3’ and ENTR-RV primer 5’- GCGACCGGCCGGTGGAT-3’. The compatible fragments were combined using HiFi DNA assembly (New England Biolabs, E2521L).

pENTR H2B mScarlet-I was created by PCR amplification of mScarlet-I from plasmid EB3- mScarlet-I (Addgene #98826, Dorus Gadella), using the primers FW 5’- ACCGGTGAATTCACCATGGTGAGCAAGGGCG-3’ and RV 5’-

GGATCCGCCTGCAGGTTACTTGTACAGCTCGTCCATG-3’. A microhomology compatible vector was created by PCR amplification of pENTR-SNAPf-H2B using primers FW 5’- CCTGCAGGCGGATCC-3’ and RV 5’- GGTGAATTCACCGGTCTGTAC-3’.

pENTR LCK-mScarlet-I was created by PCR amplification of LCK-mScarlet-I, (#98821, Addgene, Dorus Gadella), with the primers FW 5’- CCACCGGCCGGTCGCGTGAACCGTCAGATCCGCTAG-3’and RV 5’-TCGAGTGCGGCCGCTTTATCTAGATCCGGTGGATCCCG-3’. Compatible overhanging microhomologies of pENTR1a GFP N2 (FR1) were created using primers FW 5’- AGCGGCCGCACTCGAGA-3’ and RV 5’- GCGACCGGCCGGTGGAT-3’.

pENTR-GEF-H1-FLARE212-sGFP2-mScarlet-I-FRET was derived from the original GEF-H1- FLARE212 biosensor as described previously^32^ with the following modifications. The CFP-YFP FRET donor-acceptor fluorophore module was replaced with sGFP2 and mScarlet-I, retaining all the original linker regions. The entire coding sequence was synthesised as a gene block (Gene Universal) and ligated into the PUC57 plasmid. PUC57-GEF-H1-FLARE212-sGFP2-mScarlet-I- FRET was then subcloned into an entry vector by restriction digest of pENTR1a-eGFP2-N2 (FR1) and PUC57-GEF-H1-FLARE212-sGFP2-mScarlet-I-FRET with KpnI and XbaI. DNA fragments were ligated together using T4 DNA ligase. All entry plasmids were propagated in XL2-blue ultracompetent *E*. *coli* (Agilent, 200150) and sequence verified via sanger sequencing (Australian Genome Research Facility, St Lucia).

Lentiviral expression vectors were created by Gateway Gene Cloning (Invitrogen, 11791-020) with Gateway LR Clonase II (Invitrogen, 11791) into either of the following destination vectors: pLX_TRC311 (Addgene, #113668, John Doench), pLenti CMV Hygro DEST (W117-1) [9], (Addgene, #17454, Eric Campeau and Paul Kaufman), pLIX_403 (Addgene, #41395, David Root). Lentiviral plasmids were propagated in One Shot Stbl3 chemically competent *E. coli* (ThermoFisher Scientific, C737393). Plasmids were sequence verified with Sanger sequencing using vector specific primers. pLIX_403 fusion gene expression was induced using 5 μg/mL Doxycycline hyclate (Merck, D9891) for 12 h prior to any performed experiment and withdrawn during the experiment. pLenti LifeAct-eGFP BlastR (Addgene #84383, Ghassan Mouneimme).

### Lentiviral Production and Cell Transduction

Human pseudo-typed lentivirus particles were produced in low passage HEK293FT cells, co- transfected with lentiviral packaging plasmids (pCEP4-tat, pHEF-VSVG and pNHP) and lentiviral expression plasmids using Lipofectamine 2000 (Invitrogen, 11668019) as previously described^50^ with the following modifications. 24 h after transfection, the media was replaced with media supplemented with 30% v/v foetal bovine serum. Media containing high titre pseudo virus was collected at 4 h intervals after the first media exchange and continuously for 48 h thereafter and filtered through a 0.45 μm PES filter (Merck, SLHP033RS) to remove cell debris. Media containing viral particles was supplemented with 8 μg/mL Hexadimethrine bromide (Sigma-Aldrich, 107689) to transduce target cells. Infected cells were selected 48 h after transduction with antibiotic supplemented media containing either 0.1 mg/mL Hygromycin B (Roche, H3274), 5 μg/mL Blasticidin S Hydrochloride (Gibco, A1113903), 500 μg/mL Geneticin/G418 (Santacruz, 108321-42-2) or 2.5 μg/mL Puromycin dihydrochloride (Gibco, A1113803), depending on the lentiviral expression plasmid transduced. Stable cell lines were made following lentiviral transduction and selection as described previously^51^. Cell sorting was conducted using either a Beckman Coulter MoFlo Astrios EQ or BD FACSAria Fusion (SORP) (TRI *flow cytometry suite*, Translation Research Institute). Cells were sorted to isolate 2-6 populations simultaneously, using gating strategies that identified target populations based upon both the combination of fluorophores expressed and the desired level of expression for the given fluorophores.

Lentivirus-mediated shRNA was performed as above, using the lentiviral shRNA plasmid pLKO.1 expressing CLASP targeting shRNA sequences. Validated shRNA pLKO.1 target sequences were used as previously described^52^ including a non-targeting sequence that has no known target in the mammalian genome was used as a control. Transduced cells were maintained in 2.5 µg/mL Puromycin dihydrochloride (Gibco, A1113803) selection media for 5 days before experimental use. CLASP-depletion was validated by immunoblot.

Full-length 2xmNeonGreen-CLASP1 was generated using RNA isolated from 1205Lu cells. RNA was extracted using the Nucleospin RNA mini kit (Macheryl-Nagel, 740955.50). First-Strand cDNA was synthesised using the Superscript III First-Strand Synthesis System (Invitrogen, 18080051) using oligoDT primers. The CLASP1 isoform was amplified using Q5 Hot Start High-Fidelity DNA Polymerase (New England BioLabs, M0493) with gene specific primers (FW 5’- TCTGGATTTGAATTCCACTATGGAGCCTCGCATGGAG-3’ and RV 5’- TACTGCCATTAGCTGTGCGTGGAGACATCGGAGGA-3’) from synthesised cDNA. The correct target amplicon was identified by agarose gel electrophoresis and purified using a NucleoSpin gel and PCR Clean-up kit (Macheryl-Nagel, 740609). Purified DNA was ligated into the pMiniT 2.0 vector using the New England Biolabs PCR cloning kit (E1203S) and transformed into XL2-blue ultracompetent *E. coli* (Agilent, 200150). Sequences were validated using sanger sequencing with CLASP1 specific primers (CLASP1-1 FW 5’- GTGCTGCTGAGTATGATAACTTCT- 3’ and CLASP1-1 RV 5’-CTGATGACAGATGCCCCAAC-3’) designed against reference the CLASP1-NM_001142273.1 reference sequences using Snapgene software (GSL Biotech LLC). pENTR-2xmNeonGreen- CLASP1 was created by HiFi assembly of the following PCR fragments pENTR1a-GFP-N2 using the primers FW 5’– AGCGGCCGCACTCGAGA – 3’ and RV 5’- GCGACCGGCCGGTGGAT-3’, mNeonGreen-H2B-C10 using the primers FW 5’- CCACCGGCCGGTCGCATGGTGAGCAAGGGCG-3’, RV 5’- GCCCTTGCTCACCATCTTGTACAGCTCGTCCATGC-3’, mNeonGreen-H2B-C10 using the primers FW 5’-ATGGTGAGCAAGGGCG-3’, FW 5’- CATGGCGGTACCGGACTTGTACAGCTCGTCCATGCCC-3’ and pMiniT 2.0 CLASP1 using the primers FW 5’-TCCGGTACCGCCATGGAGCCTCGCATGGAG-3’ and 5’- TCGAGTGCGGCCGCTTTAGCTGTGCGTGGAGACATCG-3’. pENTR1a-2xmNeonGree- CLASP1 was recombined into pLX-TRC311 to create pLX-2xmNeonGreen-CLASP1.

2xmNeonGreen-CLASP1 stably expressing 1205Lu cells were created by first transducing cells with pLX-2xmNeonGreen-CLASP1 lentiviral particles followed by Blasticidin selection. Cells were then transduced with CLASP1 shRNA targeting the 3’ UTR of CLASP1 RNA and grown in selection in Puromycin and Blasticidin S selection for two weeks, before cell sorting based on mNeonGreen fluorescence intensity.

N and C terminally truncated CLASP1 constructs (CLASP1-Nterm, and CLASP1-Cterm, respectively) were designed by splitting full length CLASP1 between Leucine 549 and Proline 550 of the disordered region. The first 549 amino acids encoded TOG1 and TOG2, N-terminally tagged with eGFP. The amino acids 550-1470 encoded SXIP-TOG3-CLIP-ID N-terminally tagged with Halotag. Trunctated constructs were synthesised (Gene universal) and subcloned into gateway entry vectors before being recombined into the lentiviral expression plasmid pLX_TRC311 (Addgene, #11368). 1205Lu cells were then stably transduced with either pLX-eGFP-CLASP1-N (TOG1-2) or pLX-Halotag-CLASP1-C (SXIP-TOG3-CLIPID). 1205Lu-pLX-Halotag-CLASP1-C (SXIP-TOG3-CLIPID) were labelled using Halo-JF549 prior to experiments

pET28a-Halotag-CNA35 was created using InFusion HD Assembly (Takara) by replacement of the eGFP coding sequence within pET28a-eGFP-CNA35^46^ (Addgene, #61603, Maarten Merkx; primers FW 5’-TCCGGAGAATTCCACGGATCCG-3’ and RV 5’-GCTAGCCATATGGCTGCCGC-3’), with a HaloTag coding sequence from pENTR4-Halotag (Addgene, #29644, Eric Campeau; primers FW 5’- AGCCATATGGCTAGCATGGCAGAAATCGGTACTGGC-3’ and RV 5’-GTGGAATTCTCCGGAGCCGGAAATCTCGAGCGTC-3’). The construct was sequence validated before expression and purification as previously described^46^.

JF549 and JF646 Halotag and SnapTag ligands^53^ were a generous gift from Luke Lavis (Janella Research Campus, HHMI). Hoechst 33342, (Invitrogen, H1399) Annexin V 647 (Invitrogen, A23204) were used as live cell dyes.

### CRISPR Knock-in of meGFP-α-Tubulin

CRISPR-CAS9 endogenous meGFP tagging of the TUB1AB in 1205Lu cells was performed as previously described^54, 55^. To create a uniform fluorescent population of cells, cells were sorted using a MoFlo Astrios EQ cell sorter (Beckman Coulter) using a two-round sort strategy. Cells were first sorted using an enrichment mode to sort cells expressing high levels of meGFP. These enriched cells were immediately subjected to a second round of sorting, employing a purity sort mode (and gating strategy that defined a narrow window of meGFP expression), resulting in a purified population with uniform meGFP and post-sort purity of >98%.

### Antibodies, immunofluorescence and immunoblotting

Immunofluorescence was performed using protocols previously described^50, 52^. Acetylated α- tubulin (Merck, T7451), tyrosinated α-tubulin (Merck, MAB1864-I) were detected using protein specific antibodies. Goat anti-Mouse Alexa-488 (Thermo Fisher Scientific, A32723) and Goat anti-Rabbit Alex Fluor 594 (Thermo Fisher Scientific, A-11012) conjugated secondary antibodies were used for immunofluorescence. F-actin and nuclei were labelled with Alexa Fluor 647 phalloidin (Thermo Fisher Scientific, A22287) and DAPI (Invitrogen, D1306), respectively. For 3D immunofluorescence samples in Collagen, 1205Lu cells were embedded for 24 hours before being fixed with 0.25% Glutaraldehyde, 4% v/v Paraformaldehyde, 0.1% Triton-X 100 in cytoskeleton stabilisation buffer and stained with CNA35 fusion proteins -with the exception of Extended Data Fig. 1 which was DQ collagen. For immunofluorescence in microchannels, cells were first fixed in 4% Paraformaldehyde in BRB80 before removal of the PDMS device.

Immunoblotting and primary antibody dilutions were performed as previously described^50, 52^. Acetylated α-tubulin (Merck, T7451), α-tubulin (Merck, T5168), CLASP1 and CLASP2 (Absea 050801A06, and 032012H02) were detected using protein specific antibodies. Rabbit, Mouse or Rat specific HRP conjugated secondary antibodies (Cell Signalling Technology, 7074, 7076, 7077) were used to detect primary antibodies for immunoblotting and detected using Luminata Forte ECL reagent (Merck Millipore, WBLUF0100). Membranes were imaged using Bio-Rad gel documentation system, and assembled in Adobe Illustrator CC. Images were adjusted linearly to enable comparison of protein of interest to loading controls

### Tubulin Pelleting Assay

1205Lu cells (4 x10^6^) were seeded onto 10 cm tissue culture dishes and allowed to adhere for 5 h. All subsequent steps were performed on ice and with reagents chilled to 4°C. Samples were washed twice with ice cold PBS containing 4 μM Taxol. Cells were scraped and lysed into 400 μl Microtubule-stabilizing buffer containing 0.1 M PIPES, pH 6.9, 2 M glycerol, 5 mM MgCl2, 2 mM EGTA, 0.5% Triton X-100, 1x cOmplete Protease inhibitor cocktail (Merck, 04693159001) and supplemented with 4 μM Taxol to maintain microtubule stability during isolation. Cell lysates were vortexed and protein concentration was determined (Pierce BCA Protein Assay Kit, 23225). 100 µg of each sample was transferred to a new microfuge tube and centrifuged for 45 minutes at 20,000 RCF at 4°C before separating the supernatant containing monomeric tubulin from the polymerised microtubule pellet. Each pellet was then resuspended into 100 µl of 1 x Laemmli buffer containing 0.5 % w/v Bromophenol blue, 0.5 M Dithiothreitol, 5% w/v SDS in 0.5 M Tris- Cl pH 6.8 and incubated at 95°C for 5 minutes. 20 µg of each cell protein lysate sample was separated using a hand-cast 5-15% gradient Tris glycine gel (Bio-Rad, Protocol Bulletin 6201) using a custom made 25-well comb.

### Microchannel Fabrication

Fabrication of moulds for microchannel experiments was performed at the Australian National Fabrication Facility - Queensland at the Australian Institute for Bioengineering and Nanotechnology, The University of Queensland. Fabrication was performed in either a class 1000 or 10,000 clean room with generous assistance from ANFF-Q professional staff.

### Chrome Mask Printing

Microchannel designs were created using L-Edit (Tanner Tools) or AutoCAD 2021 and printed using a Heidelberg uPG 101 using the high-resolution print head (> 2 µm features). Designs were etched onto 5-inch square soda-lime chrome masks. After chrome photomask etching, each mask was subjected to one cycle of post baking on a dry hot plate at 95°C for one minute. Each chrome mask was developed in AZ726 developer (MicroChemicals) for 45 seconds with gentle agitation, then rinsed thoroughly with de-ionised water and dried with nitrogen gas. Chrome etching of each photomask was performed using Chromium etchant (Merck, 651826). Timing varied between 1- 10 minutes, notably the termination of the process was finalised after the following steps were observed: 1. Chrome appearance of features 2. Chrome disappearance 3. Feature transparency at locations originally seen in chrome in step 1. Once progression through steps was visualised, each mask was rinsed with deionised water and dried with nitrogen gas. To complete chrome photomask development, each mask was submerged in acetone and sonicated in an ultrasonic cleaner at 30% power at 75°C. Post sonication, each mask was washed several times with acetone, then placed into another beaker filled with 100% isopropanol and sonicated under the same conditions. After sonication, masks were washed one final time with 100% isopropanol and each mask was gently wiped clean with a low lint towel wetted with 100% isopropanol.

### SU-8 2000 Mould Fabrication and Silanization

To ensure SU-8 2000 adhesion to silicon wafers, 5-inch round silicon wafers were etched using 48% hydrofluoric acid (Sigma, 695068). The etched wafer was then washed with water, dried using a nitrogen gas and used immediately for SU-8 2000 (Kayaku Advanced Materials, MicroChem) spin coating. To achieve a ∼4 µm thickness, SU-8 2000 was spin coated as per manufacturer’s instructions. The subsequent SU-8 2000 was then soft baked as per manufacturer’s instructions. Chrome masks were then overlaid onto SU-8 2000 spin-coated silicon wafers. The resulting chrome mask SU-8 2000 sandwich was exposed on an EVG620 (EV Group) mask aligner using ∼360 nm UV light, exposure time and power was tailored depending on mask thickness (typically UV dose ranged from 120 mJ/cm^2^ – 140 mJ/cm^2^). SU-8 2000 regions exposed to UV were then developed using manufacturer instructions.

After SU-8 2000 moulds were developed, each mask was silansed to create an anti-stiction layer for easy PDMS demoulding from SU-8 2000 silicon wafers. Silanization was performed by applying two drops of Trichloro (1*H*,1*H*,2*H*,2*H*-perfluorooctyl) silane (Merck, 44893) onto a sheet of aluminium foil, then placed into a vacuum desiccator alongside the SU-8 2000 mould. Air was evacuated using a vacuum tap for 5 minutes, after which the vacuum line was clamped. The vacuum was held for an hour or until the silane had completely vaporised. Successful silanisation of the mask was observed by a faint opaque layer on the surface of the SU-8 2000 mould. This step was performed once for every new mould developed.

### Polydimethylsiloxane (PDMS) Casting

PDMS (Slygard 184, Dow Corning) was mixed at a ratio of 10:1 (w/w) base to catalyst-cross linker and poured on top of SU-8 2000 moulds, then baked at 60°C for 2 hours. Seeding ports were created using a 3 mm biopsy punch (T983-30, ProSciTech). PDMS microchannels were permanently bonded to glass using a Corona discharge tool. Microchannel devices were functionalised with 50 μg/mL collagen I in PBS and incubated overnight at 37°C before replacing with media. To perform experiments, between 1.0 e^4^-5.0 e^4^ cells were resuspended in 10 μl and seeded into each 3 mm port. Cells were allowed to attach for 6-12 hours before imaging.

### Axial Confinement of Cells

*Confinement device.* Confinement was controlled using a modified confinement piston^56^. Briefly, to induce and maintain compression over 24 hours, a custom designed PDMS piston was cast from a series of machined aluminium rings to create a PDMS suction cup mould. PDMS was then cast from the mould and connected to a 60 mL syringe (BD Bioscience) via silicone tubing using Luer lock connectors. To connect the silicone tubing to the PDMS piston, a blunt 18-gauge needle was bent 90 degrees and the plastic connector removed. The needle was then inserted into a hole in the side wall of the PDMS, using a 1 mm biopsy punch, lubricated with silicon grease at both ends. The alternate end of the silicone tubing was connected to a barbed female Luer connector and then connected to a syringe. To induce axial confinement, a generous coating of silicone grease was first applied to the contact surface of the PDMS suction cup to create an airtight seal. Syringes plungers were displaced backwards by a 2-5 mL graduation and negative pressure was locked in place using a 60 cc Shippert syringe snap lock (SNAP60, Precise Medical Supplies).

*Axial confinement assays:* Cells were seeded onto 35 mm glass bottom dishes (Cellvis, D35-20- 1.5H) coated with poly-d-lysine (0.1 mg/mL). Confinement height was controlled using a coverslip containing 3 µm height PDMS micropillars passivated with non-adhesive pLL-PEG (SuSoS). Microtubule stability was assessed via axial confinement of 1205Lu cells endogenously labelled for meGFP-α-Tubulin (CRISPR) stably co-expressing pLenti-CMV-blast-snapf-H2b (labelled with SnapJF549) in control (non-targeting) and CLASP shRNA transduced cells. Membrane rupture was assessed using intracellular propidium iodide influx during confinement experiments. Total cell nuclei were counterstained with 1.3 µM Hoechst 33342 (Thermo fisher scientific) diluted in PBS and stained for 10 minutes, followed by five washes with PBS (5 minutes per wash). Cells were incubated in 50:50 DMEM:F12 media, 10% foetal bovine serum (Gibco, 10100147), 0.1 mM MEM non-essential amino acids (Gibco, 11140050) and 100 i.u. mL−1 penicillin, 100 μg mL−1 streptomycin (Gibco, 15070063) containing 1 µg/mL Propidium iodide (Sigma, P4170) to identify loss of membrane integrity, and 50 mM HEPES to buffer media pH in the absence of CO_2_.

### Microscopy, Image Processing and Data Analysis

Fixed and live imaging (unless otherwise stated) was performed on a custom-built spinning disc confocal microscope as previously described^51^ or an Andor Dragonfly spinning disc confocal equipped with dual Andor Zyla 4.2 sCMOS cameras, controlled by Fusion Software (Andor). 3D- SIM was performed using a GE-OMX-V4 Blaze (Applied Precision, GE Healthcare). LSFM was performed on a custom built high numerical aperture variant of Axially Swept Light Sheet Microscope without utilising remote focusing or rolling shutter readout, details of this system are published elsewhere^12, 57^. Brightfield and epifluorescence live cell microscopy was performed on a Nikon Ti-E inverted microscopy equipped with a Hammamatsu Flash 4.0 sCMOS camera, controlled by NIS Elements software (Nikon) or an Olympus IX81 live cell microscope equipped with a Hamamatsu Orca Flash 2.8 camera. Subcellular fluorescent protein dynamics were imaged at 37°C, supplemented with 5% CO_2_ (Okolab Cage Incubator Gas Chamber, Bold Line Gas Controller operated by OKO-TOUCH).

Raw images were processed into images and movies in Fiji and NIS elements (Nikon). Tiff stacks were bleach corrected and sharpened using the unsharp mask tool for visualisation purposes only. Quantification was performed only on raw 14-16-bit images. Black-magenta image overlays were generated as described previously^52^. Figures were assembled using Adobe Illustrator CC (Adobe). Statistical analysis was performed in GraphPad Prism 9.0, R Studio, Excel or PlotTwist^58^. For power in statistics, we assumed unequal variance and non-normal distribution. A Mann-Whitney U test was used for statistical comparison between two groups. A Kruskal-Wallis One-way ANOVA test with a Dunn’s correction for false discovery was used for comparisons of two or more groups. Reproducibility was determined based on at least three individual repeats. No predetermined sample sizes were arranged during experimentation.

*Object tracking*. Cells were embedded in collagen gels (described above) then recovered in a tissue culture incubator for an hour prior to imaging for 24 h. Images were acquired every 5 min. Cells were tracked based on mCherry-H2B fluorescence using Trackmate^59, 60^. XY tracking data was used to create representative spider-plots, as previously described^50^. CLASP1 tracking was performed using the manual tracking function in Fiji.

*Cell viability in microchannels.* Nuclei morphology was used to determine cell viability within constriction microchannels. Cells death was deemed to occur once nuclear condensation and fragmentation appeared in the earliest frame (Hoechst 33342 stained or Snaptag-H2B fluorescence).

*Microtubule windowing analysis.* Windowing analysis was performed in MATLAB as previously described^61^. Images of cells expressing meGFP-α-tubulin were first cropped in half along the long axis of the cell in the direction of cell migration. Sampling windows were held constant throughout all frames. Only the first windows adjacent to the defined cell membrane edge was quantitated to derive fluorescence intensity values from microtubules which experienced the most direct mechanical force. Window values were averaged with surrounding nearest neighbour values and normalised to the first sampled window and shown as fold-change. Windows containing NaN values were excluded and replaced with 0. Heatmaps were plotted in R studio using ggplot2^62^.

*CLASP dynamics in constriction microchannels.* 1205Lu cells stably overexpressing 2xmNeongreen-CLASP1 and plKO.1-shRNA targeting endogenous CLASP1 (#31) were seeded into constriction microchannels and allowed to migrate for 24 hours prior to imaging. Time-lapse imaging of CLASP1 dynamics was acquired using an inverted Zeiss LSM 880 with the fast Airy scan mode at 1 second intervals for a total duration of 60 seconds.

*Microtubule abundance during axial confinement.* Image stacks were thresholded to mask pixels of polymerised microtubules without detecting unpolymerised cytoplasmic tubulin. Binary masks were generated from thresholded images and used to measure the total surface area over time.

*3D kymograph timeseries.* 3D kymographs were generated in Fiji by firstly removing backgrounds of image series surrounding the pixels of interest. To do this, background binary mask series were generated by thresholding, then multiplied against the original image series. Time-series were then reordered by swapping Z and Time dimensions. Pseudo-Z-series were then 3D projected.

*Propidium Iodide cell viability analysis.* 3-5 representative images per time point were acquired, taken using an inverted epifluorescent EvoS microscope. Propidium iodide positive nuclei and total nuclei were counted manually using the multi-point tool in Fiji. Propidium iodide influx frequency was calculated by dividing total propidium iodide positive nuclei by total cell number and displayed as a percentage as a function of time in Excel (Microsoft). Statistical significance was determined at individual time points using an unpaired Mann-Whitney test assuming non-gaussian distribution of data using GraphPad Prism (9.4.1).

*Microtubule to cytoplasmic intensity ratio analysis.* 1205Lu cells stably expressing pLX-eGFP- CLASP1-N were axially confined as above. To determine microtubule binding, a circular ROI the size of an individual microtubule was placed at locations corresponding to the microtubule polymer or cytoplasm, respectively. A total of 15 individual ROI locations we measured per time point, per condtion to measure the mean intensity over time. Intensity measurements were background subtracted and displayed as a ratio of microtubule/cytoplasmic intensity as a function of time.

*Nuclear approach angle, morphometrics, passage time, rupture and curvature.* Nuclear approach angles were measured using the ‘angle tool’ in Fiji. Angles were created using the first frame where individual nuclei were seen to contact constrictions. Angles were measured by drawing a centre line between the two pillars of constriction in the direction of migration, the second line was drawn to cover the long axis of the nucleus. The angle between the two lines was determined as the nuclear approach angle. Nuclear morphology was quantitated using fluorescent images of Janelia Fluor 549/646 covalent linked Snaptag-H2B nuclei. Images were first gaussian blurred using a sigma of 2.0 then binarised, any holes present within binarised nuclear shapes were filled using the ‘Fill Holes’ function before using the ‘Erode’ function. Binary masks were used to derive nuclear circularity measurements and XY positions over time. Extracted values were then used to calculate displacement and instantaneous speed. Duration of nuclear passage time was determined by counting frames between from nuclear constriction pillar first contact to the last frame in which nuclei had completely cleared constriction channels. Nuclear rupture was quantitated based on changes in 3xNLS-mScarlet-I-NLS signal intensity. The magic wand tool (legacy mode) was used to create ROIs which encompassed at least 90% of nuclei as cells progressed through constrictions. Fluorescence intensity measurements were normalised to the first measured frame to determine fold-change over time. A cut-off of 0.04-fold reduction was used to determine the presence of nuclear rupture events. Nuclear circularity and nuclear rupture measurements were plotted as a function of normalised time, individual dynamic traces were normalised to each other by aligning the lowest circularity or fold-change values in each trace to timepoint 0. Nuclear curvature was measured using Kappa (*κ*)^63^ in Fiji at the front and rear nuclear contours that were not in direct contact with constriction pillars.

*Bleb analysis.* To visualise cortex (F-actin) and plasma membrane dynamics, cells were dual-transduced with pLenti6.2 eGFP-Lifeact and pLX-LCK-mScarlet-I. Cells were allowed to migrate into channels and imaging acquisition was nested at hourly intervals over a total duration of 24 h, each nested acquisition was acquired at 1 Hz for 2 mins. Kymographic curves of bleb traces were created in Fiji using a line that only spanned entire blebs from protrusion to retraction. Kymographs were used to measure: bleb formation time (the X-Euclidian-distance from cortical F-actin signal loss - marking cortex rupture- to the maximal membrane bleb height), bleb size (the maximal Y protrusion height from cortex to the peak distention of each bleb) and bleb retraction time (the X- Euclidian distance starting from the base of the maximum curve peak to complete cortex recovery). Individual bleb retraction or formation speeds were calculated using the formula: 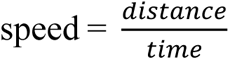.

*Microtubule-Contractility Axis Localisation Dynamics.* Quantitation of Microtubule, RhoA and Myosin dynamics during constriction was performed using a 10-point thickness segmented line drawn over the rear and front perimeter of the cell. Nuclei served as demarcation points to create front and rear compartments. Measurements from three specific stages of constricted migration were used. Early constriction – where the cytoplasm but not the nucleus touched constriction pillars, mid constriction - when the nucleus exhibited a ‘figure eight’ shape and was under peak constriction between pillars, and late constriction - when the nucleus had completely exited and cleared constriction pillars. Individual measurements were background subtracted and mean +/- SEM was graphed.

*GEF-H1 FRET*. Ratiometric GEF-H1 FRET images were created in Fiji. sGFP2 timeseries images were first binarised then multiplied, using the ‘Image calculator’ function, against individual FRET and donor channels to remove spurious FRET signal from the background. Background masked FRET channels were then divided by donor channels to produce ratiometric FRET/Donor images. To quantitate GEF-H1 activation, a polygon ROI was drawn around the rear and front of the cell cytoplasm during the previously defined phases of constricted migration to read out FRET/Donor ratios.

## Supporting information

Supplementary Video 1: Microtubules form a dynamic cage in cells moving through complex 3D environments

Supplementary Video 6: Axial confinement induces CLASP re-localisation.

Supplementary Video 7: CLASP-depletion results in cortical blebbing.

Supplementary Video 8: CLASP-depletion disrupts the proximal enrichment of myosin during confined migration.

Supplementary Video 9: Timed GEF-H1 activation upon microtubule depolymerisation precedes nuclear transmigration.

Supplementary Video 10: A timed Rho-burst at the cell-rear progresses nuclear transmigration during confined migration.

Supplementary Video 11: Simulation of nuclear passage and the microtubule cushion.

Supplementary Video 2: Contrasting microtubule organisation in 2D vs. 3D migrating cells.

Supplementary Video 3: A microtubule cage assembles in cells undergoing constricted migration.

Supplementary Video 4: CLASP depletion results in cell rupture during confined migration.

Supplementary Video 5: Axial confinement results in microtubule depolymerisation in CLASP-depleted cells.

Mathematical Model

## Extended Data Figure Legends

**Extended Data Fig. 1.**
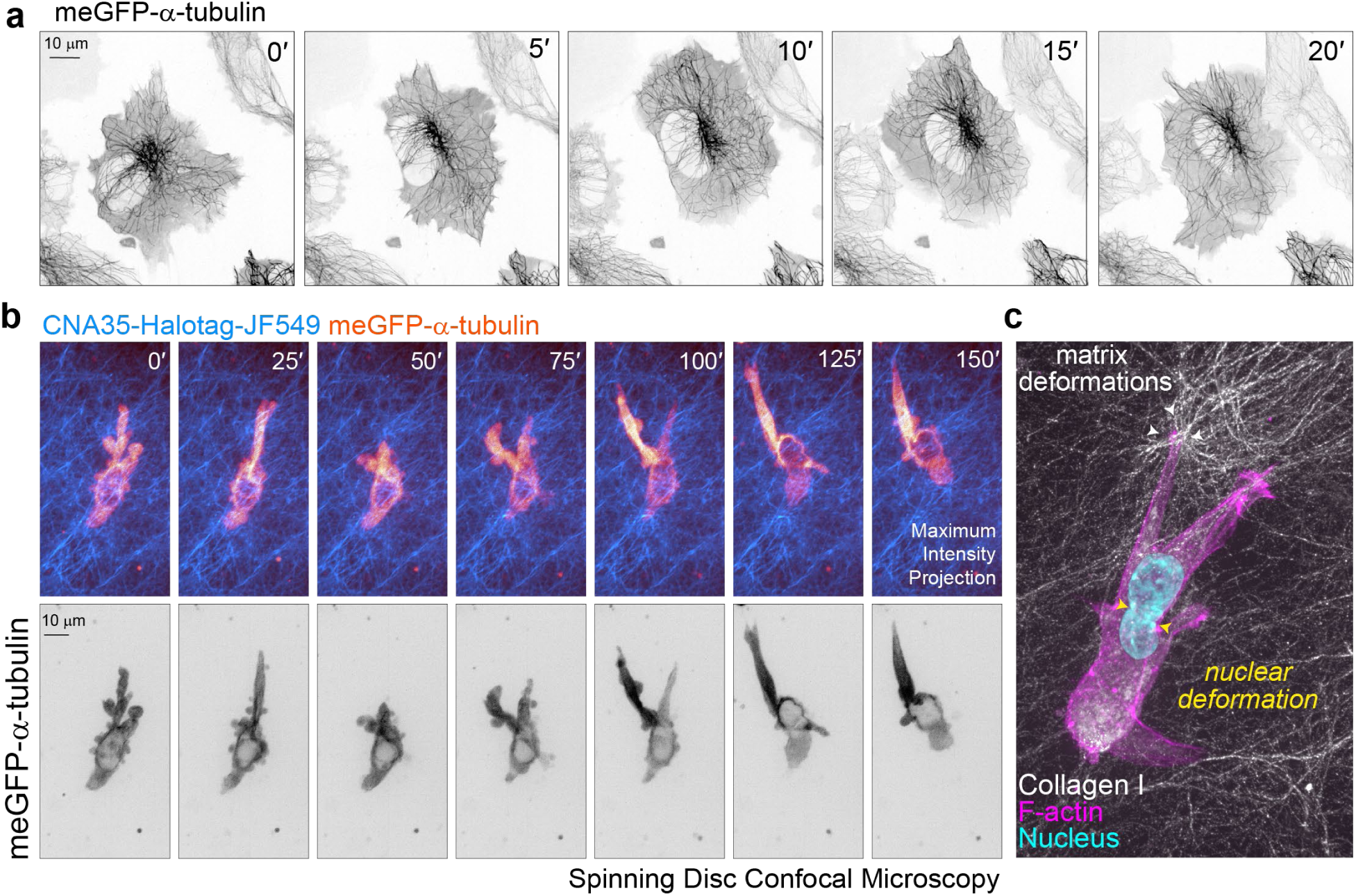
Microtubule morphology in 2D and 3D collagen hydrogel models. (**a**) 1205Lu melanoma cell CRISPR edited to express meGFP-α-tubulin migrating in 2D. LUT inverted to enhance visibility of microtubule organization. (**b**) 1205Lu-meGFP-α-tubulin (orange heatmap) expressing cell migrating through a CNA35-Halo JF549 labelled collagen I hydrogel (2.5 mg/mL; cyan heatmap) imaged by spinning disc confocal microscopy (SDCM). Visible distention of the cell body and nucleus can be observed where microtubules are seen to outline. (**c**) 1205Lu melanoma cell invading through a collagen hydrogel (DQ Collagen; white) labelled for F-actin (phalloidin, magenta) and nucleus (DAPI, cyan). Collagen aligns at cell-matrix attachment points (white arrowheads). The nucleus deforms as the cell invades (yellow arrowheads).

**Extended Data Fig. 2.**
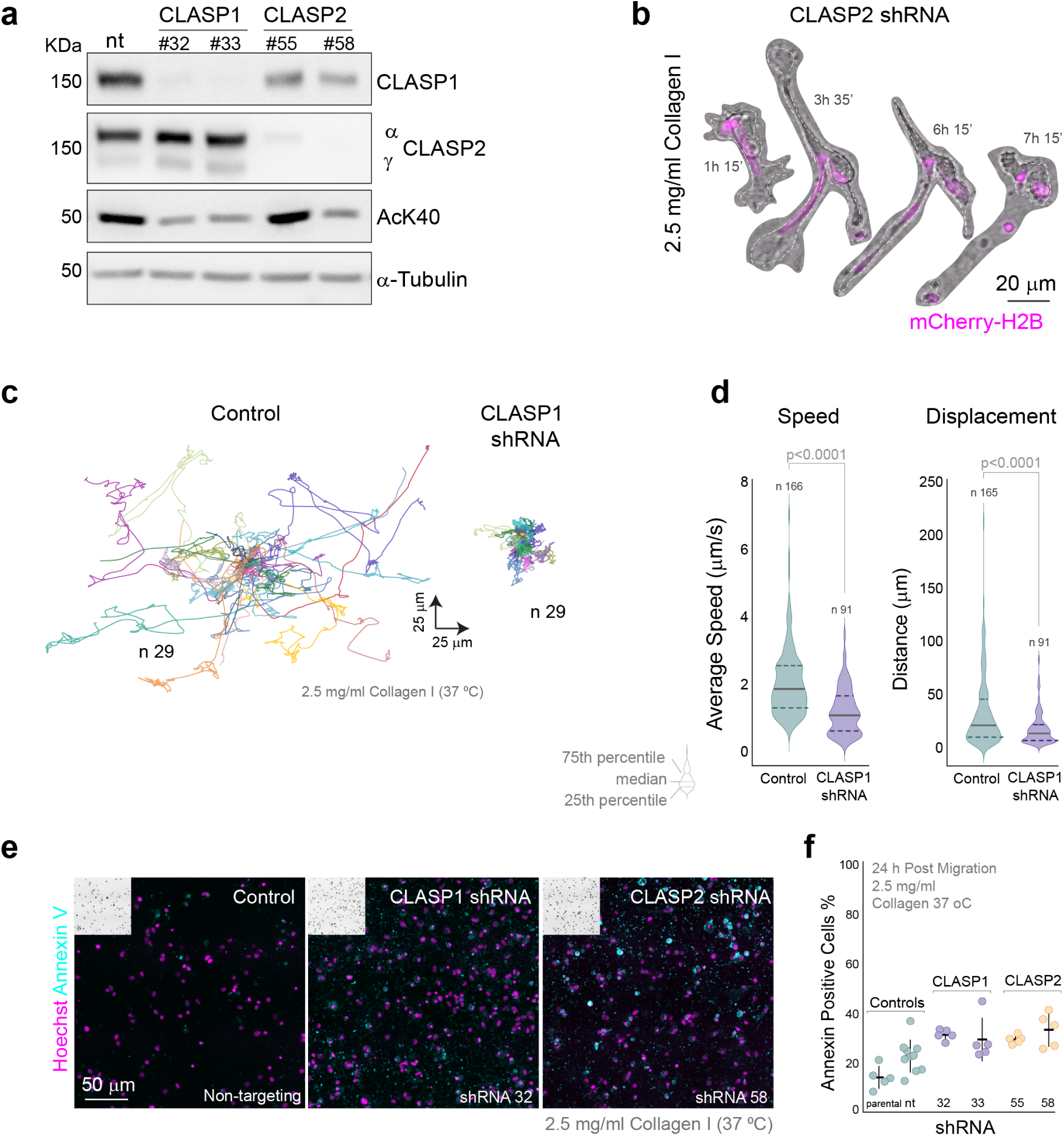
Validation of CLASP migration phenotype in 3D collagen hydrogels. (**a**) Immunoblot of lysates from 1205Lu melanoma cells expressing different shRNA constructs after seven days of puromycin selection. Tubulin was used as a loading control. Blots were probed with paralog specific antibodies to either CLASP1 or CLASP2. (**b**) Representative frames of time-lapse images of 1205Lu melanoma cells expressing mCherry-H2B (nuclei, magenta) depleted of CLASP2, embedded in a collagen I hydrogel (2.5 mg/mL). (**c**) Cell migration tracks (spider plots) of control (non-targeting) and CLASP-depleted (CLASP1 shRNA) cells. (**d**) Speed and Displacement measurements cells migrating in collagen hydrogels. (**e**) Immunofluorescence of control (non-targeting) and CLASP1- and CLASP2 depleted 1205Lu melanoma cells labelled for annexin V (cyan) and nuclei (Hoechst; magenta). (**f**) Quantitation of percentage of Annexin v positive cells in control and CLASP-depleted cells.

**Extended Data Fig. 3.**
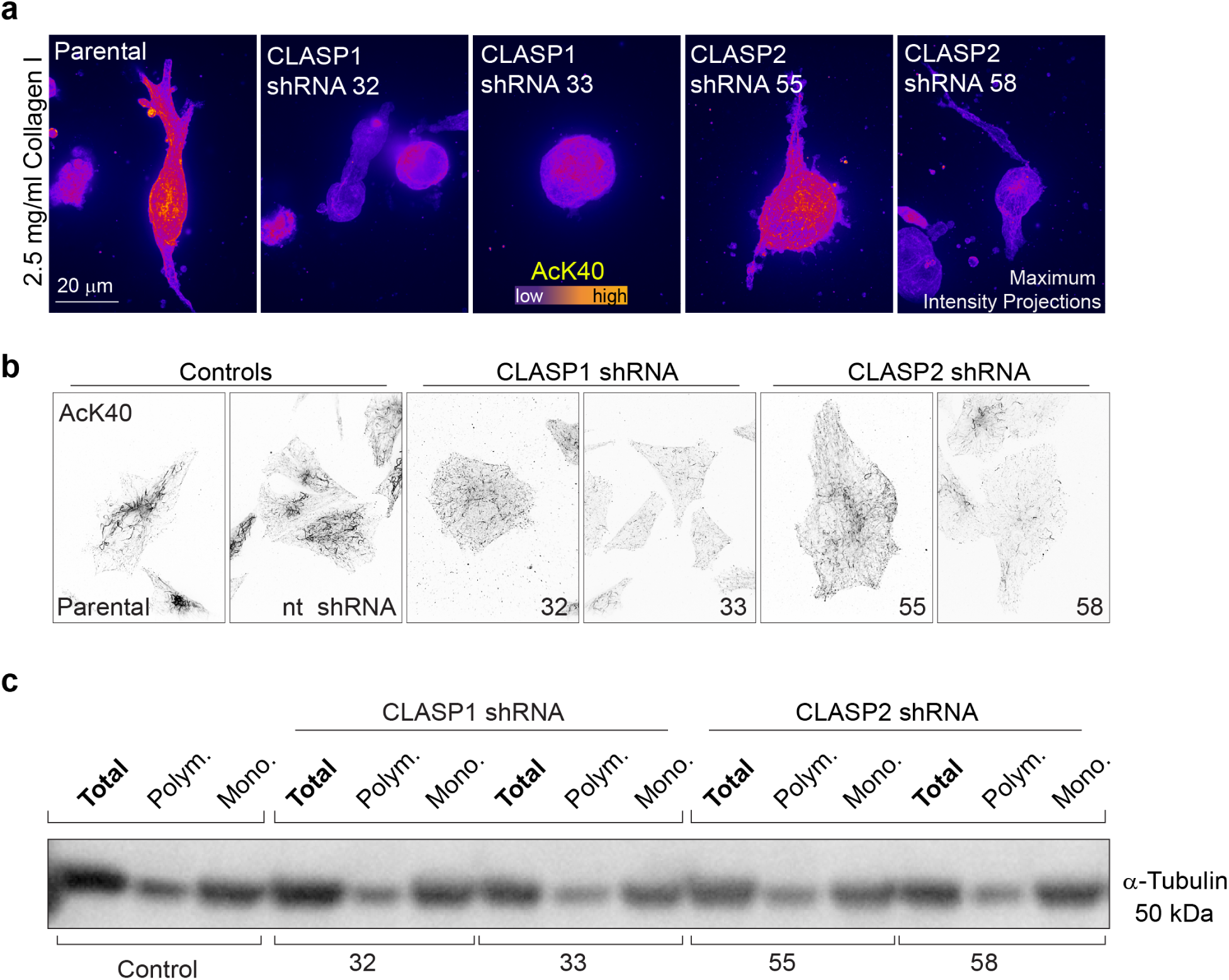
Tubulin acetylation is decreased in CLASP-depleted cells. (**a**) Control (parental), and alternative CLASP1 and 2 hairpins from figure 1F (shRNA #32, #33 and #55, 58, respectively), 1205Lu melanoma cells embedded in collagen I hydrogels (collagen, 2.5 mg/mL) labelled for acetylated α-tubulin (AcK40). AcK40 fluorescence intensity displayed as a LUT heatmap; purple indicating low with yellow-red indicating high. (**b**) Immunofluorescence of control (parental, nt; non-targeting), CLASP1-(shRNA#32, #33) and CLASP2-depleted (#55, #58) 1205Lu melanoma cells labelled for acetylated α-tubulin (AcK40, contrast inverted). (**c**) Immunoblot of tubulin pelleting assay lysates of 1205Lu melanoma cells expressing non-targeting (control), CLASP1 or 2 shRNA constructs. CLASP-depletion does not alter the levels of polymerised microtubules.

**Extended Data Fig. 4.**
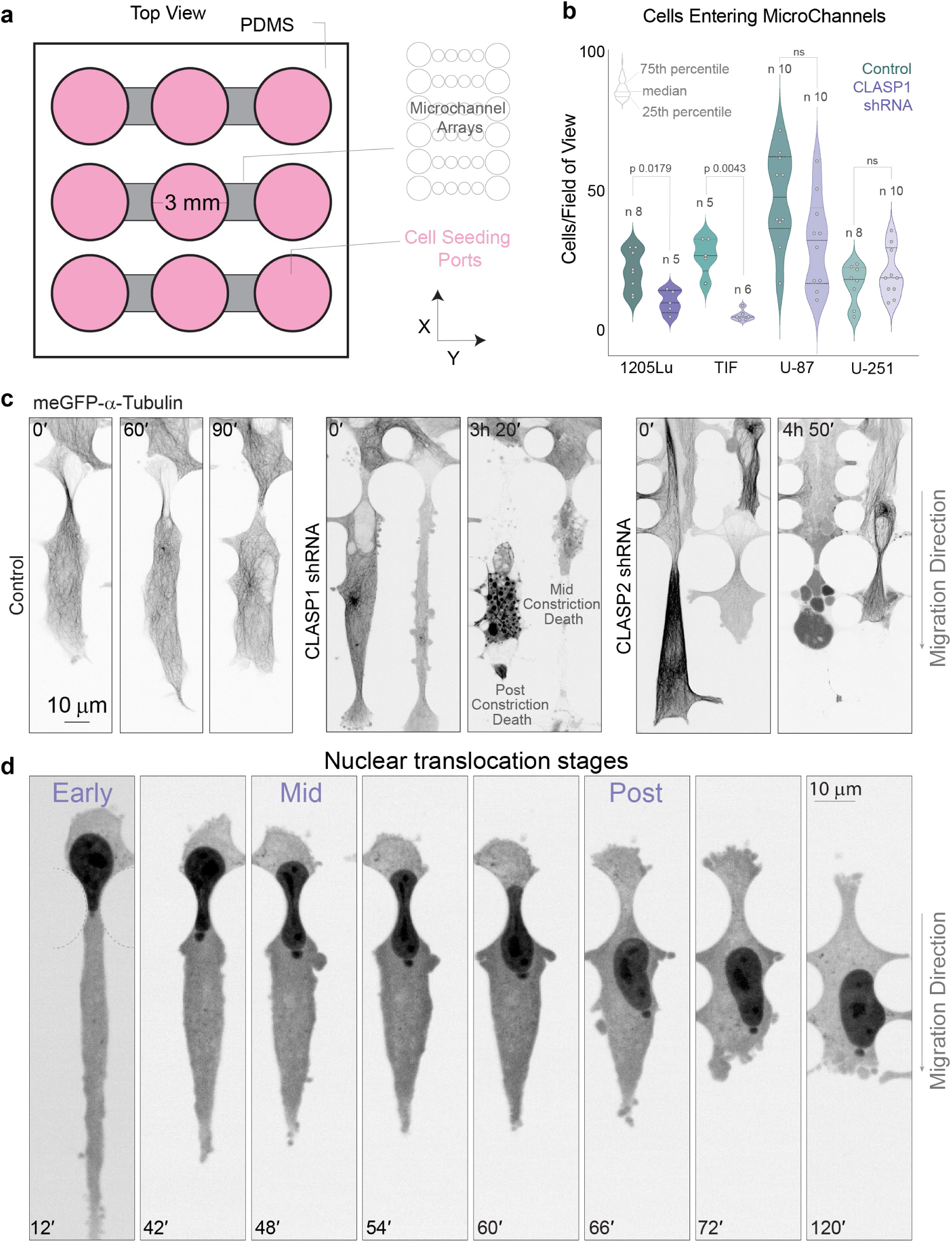
Validation of CLASP Migration Phenotype in PDMS constriction channels. (**a**) Simplified graphical schematic of constriction microchannel design. (**c**) CLASP depletion decreases the number of cells entering constriction microchannels (12 h of live imaging). Non-targeting; teal shades and CLASP1 shRNA; lilac shades), 1205Lu melanoma, TIF telomerase-immortalised normal fibroblasts, U87 and U-251 glioma. (**c**) LUT inverted timelapse sequences (spinning disc confocal microscopy) of control (non-targeting), CLASP1 and CLASP2-depleted 1205Lu cells endogenously expressing tagged microtubules (meGFP-α-tubulin). (**c**) Example of a nuclear translocation stages- early, mid and post constriction phases. Nuclear bleb depicts nuclear herniation event as a result of constriction.

**Extended Data Fig. 5.**
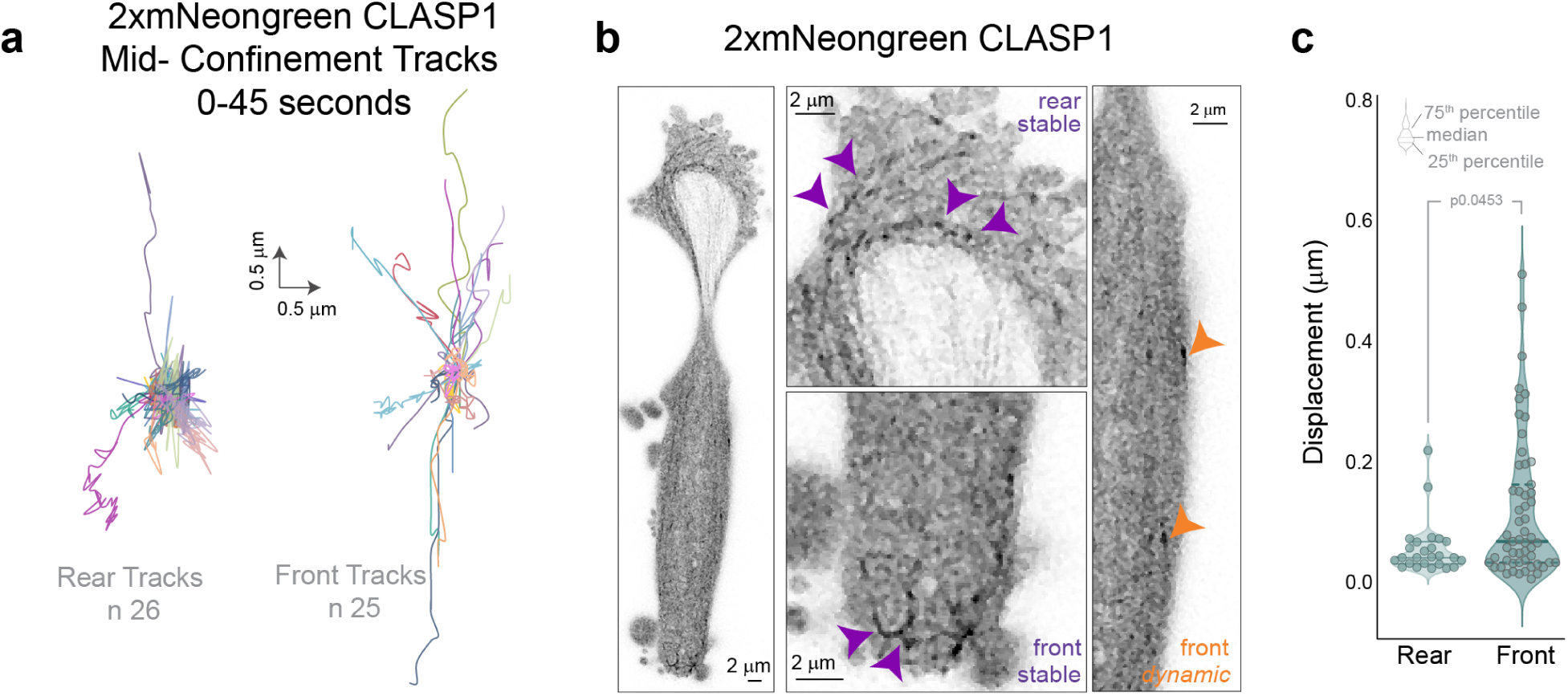
CLASP1 exhibits localised spatio-temporal dynamics. (**a**) Spider plots of CLASP1 (2xmNeonGreen-CLASP1) tracks in front and rear cell compartments during mid-nuclear occlusion. 45 seconds duration. (**b**) 1205Lu cell expressing 2xmNeonGreen-CLASP1 migrating through a microchannel construction. Magnified regions displaying CLASP dynamics in rear and front compartments. (**c**) CLASP1 comet displacement in rear and front cellular compartments during constriction.

**Extended Data Fig. 6.**
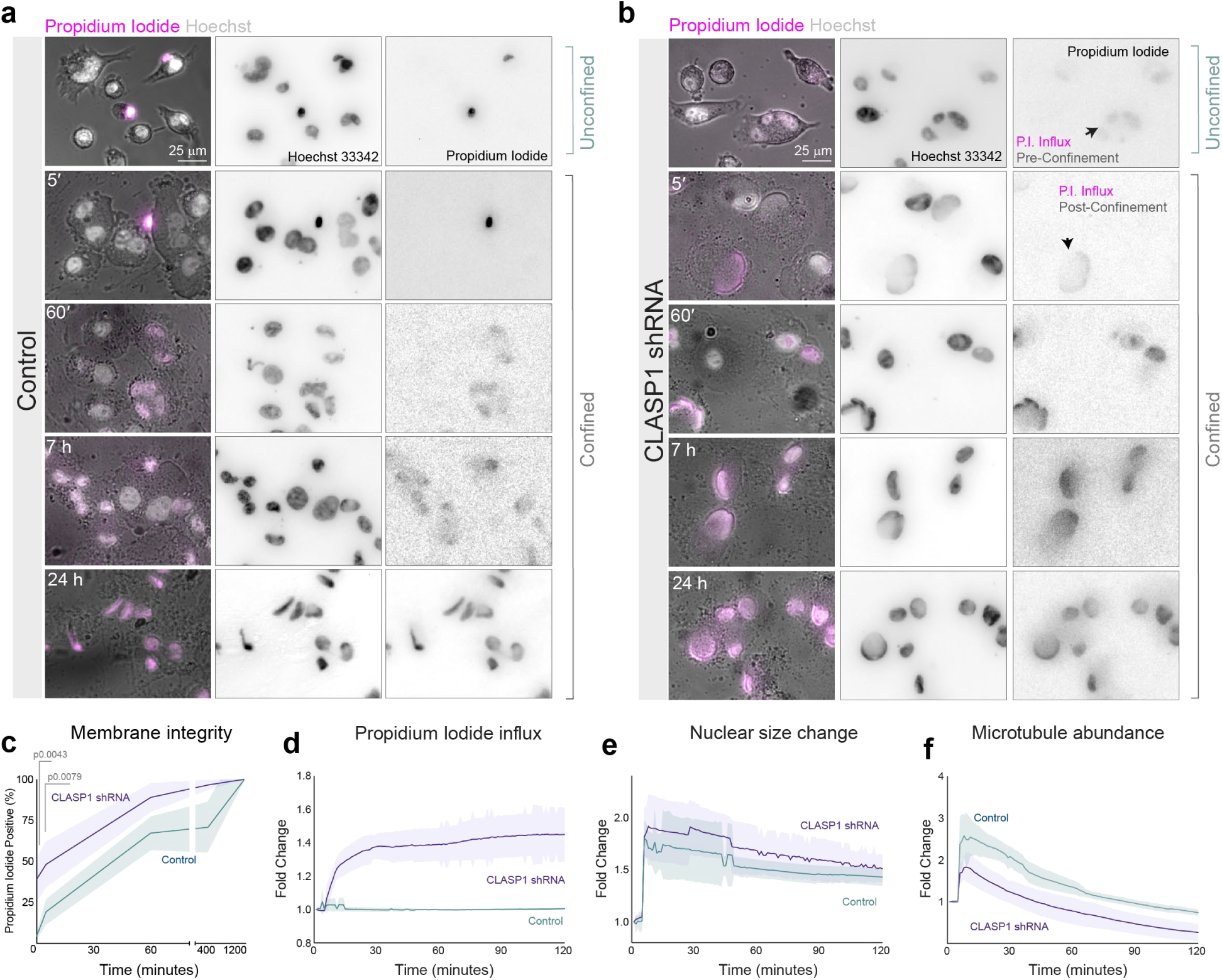
CLASP Depletion Results in Increased Membrane Rupture and Propidium Iodide influx. Representative images of (**a**) control (non-targeting) and (**b**) CLASP1- depleted (CLASP1-shRNA) 1205Lu cells subjected to axial confinement (3-micron height). Images were acquired at indicated time intervals. An increase in Propidium iodide (P.I. magenta) influx in cells counterstained with cell permeable Hoechst (nuclei, grey) was used to detect a loss of membrane integrity. (**c**) Quantitation of membrane integrity as measured by propidium iodide positive nuclei. (**d**) Quantitation of P.I. influx during axial confinement of control (non-targeting) or CLASP1 depleted (CLASP1 shRNA) 1205Lu cells endogenously expressing meGFP-α-tubulin and the nuclear marker Snapf-H2B-(Snap-JF646). (**e**). Fold change in nuclear size (area). (**f**) Microtubule stability during axial confinement as measured by fold change in microtubule area (abundance). (Relates Supplementary video 5).

**Extended Data Fig. 7.**
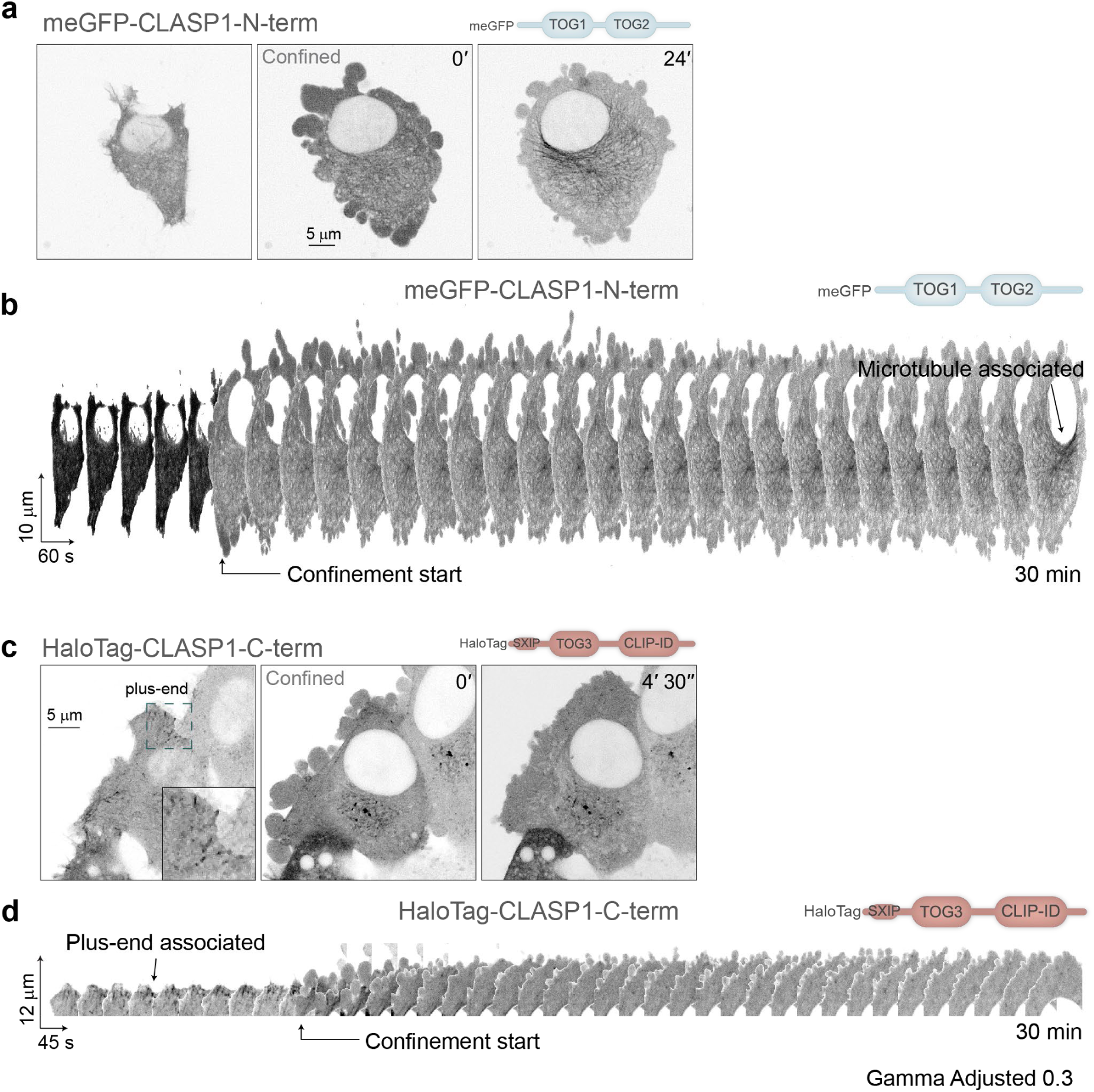
CLASP truncation mutants differentially localise on microtubules in response to axial confinement. (**a**) representative timelapse images of a 1205Lu cell stably overexpressing meGFP-tagged N-terminal fragment of CLASP1 (meGFP-CLASP1 N-term; TOG1 and TOG 2) pre- and post-axial confinement (3 microns). In the absence of confinement, meGFP-CLASP1-N-term localises to the cytoplasm. Induction of confinement results in an association with microtubules, progressively increasing overtime (24’). (**b**). 3D kymographs of the cell in a. (**c**). 1205Lu cells stably overexpressing HaloTag-tagged C-terminal fragment of CLASP1 (HaloTag-CLASP1-C-term; SxIP-TOG3-CLIPID), labelled with Halo-JF549. HaloTag-CLASP1- C-term tracks microtubule plus-ends irrespective of of axial confinement (pre-confinement 0’, and post-confinement 4’30”). (**d**) 3D kymographs of cell in c. Images were gamma adjusted (0.3) to increase contrast of microtubule structures.

**Extended Data Fig. 8.**
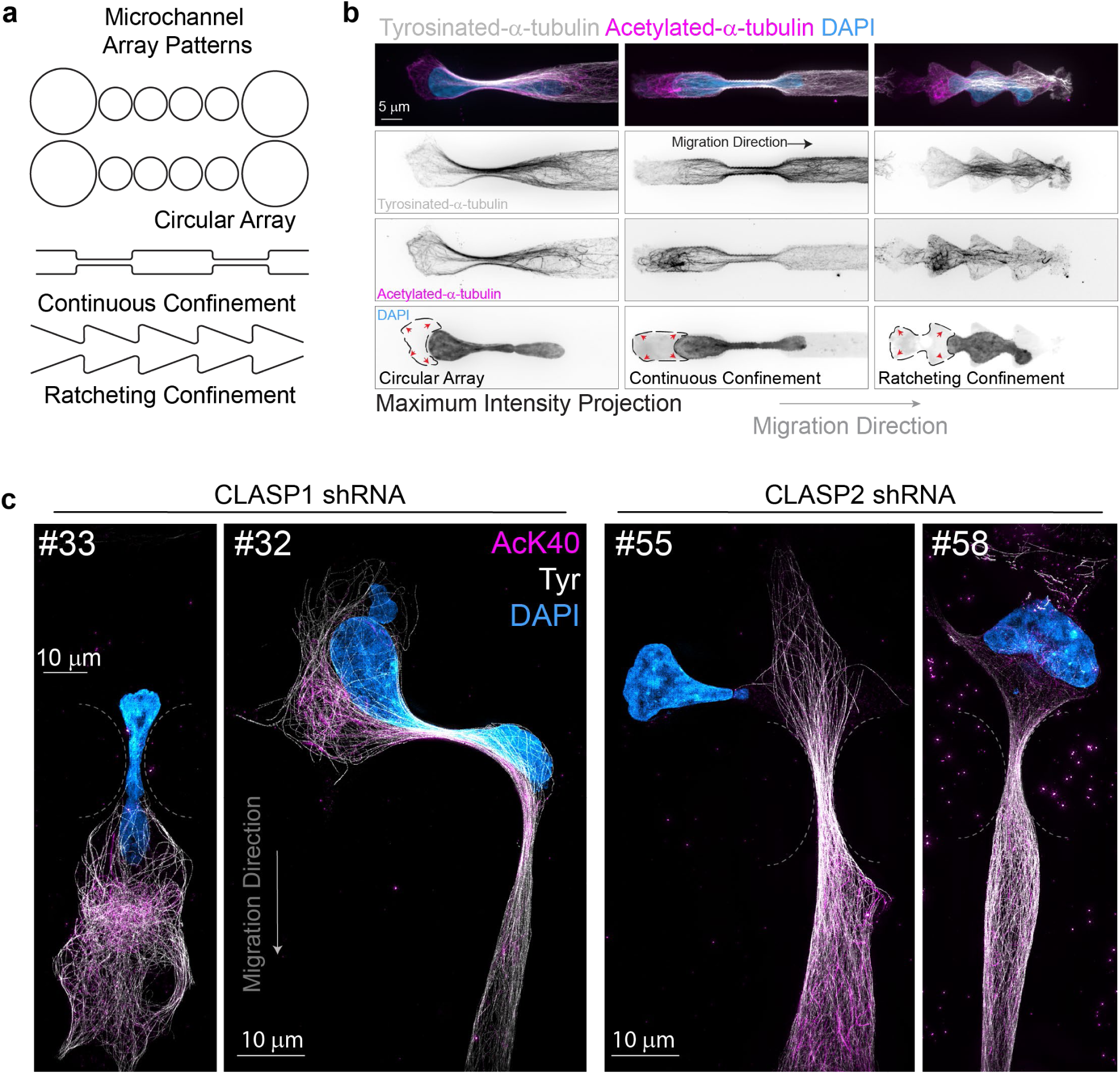
Microtubule acetylation patterns in cells in confinement. (**a**) Simplified graphical schematic of constriction microchannel patterns; Circular, continuous and ratcheting. (**b**) Immunofluorescence of 1205Lu melanoma cells labelled for acetylated α-tubulin (AcK40, magenta), tyrosinated a-tubulin (white) and nuclei (DAPI, cyan) navigating through indicated microchannel designs. Scale bar 5 µm. Images are displayed as maximum intensity projections of z-stacks. Single channels are displayed as contrast inverted images, demonstrating regardless of design, acetylated-α-tubulin forms a polarised network that contrasts tyrosinated-α-tubulin. A rear cytoplasmic compartment can be seen which flanks the rear of the nucleus (dotted line), predominantly filled with stable acetylated tubulin (red arrow heads) forming the microtubule cushion. (**c**) 3D-Structured Illumination microscopy of acetylated tubulin (AcK40) and tyrosinated α-tubulin (white) immunolabelled 1205Lu cells, nuclei labelled with DAPI, undergoing confined migration. Cells are expressing alternative CLASP1 and 2 hairpins from Fig. **2b**.

**Extended Data Fig. 9.**
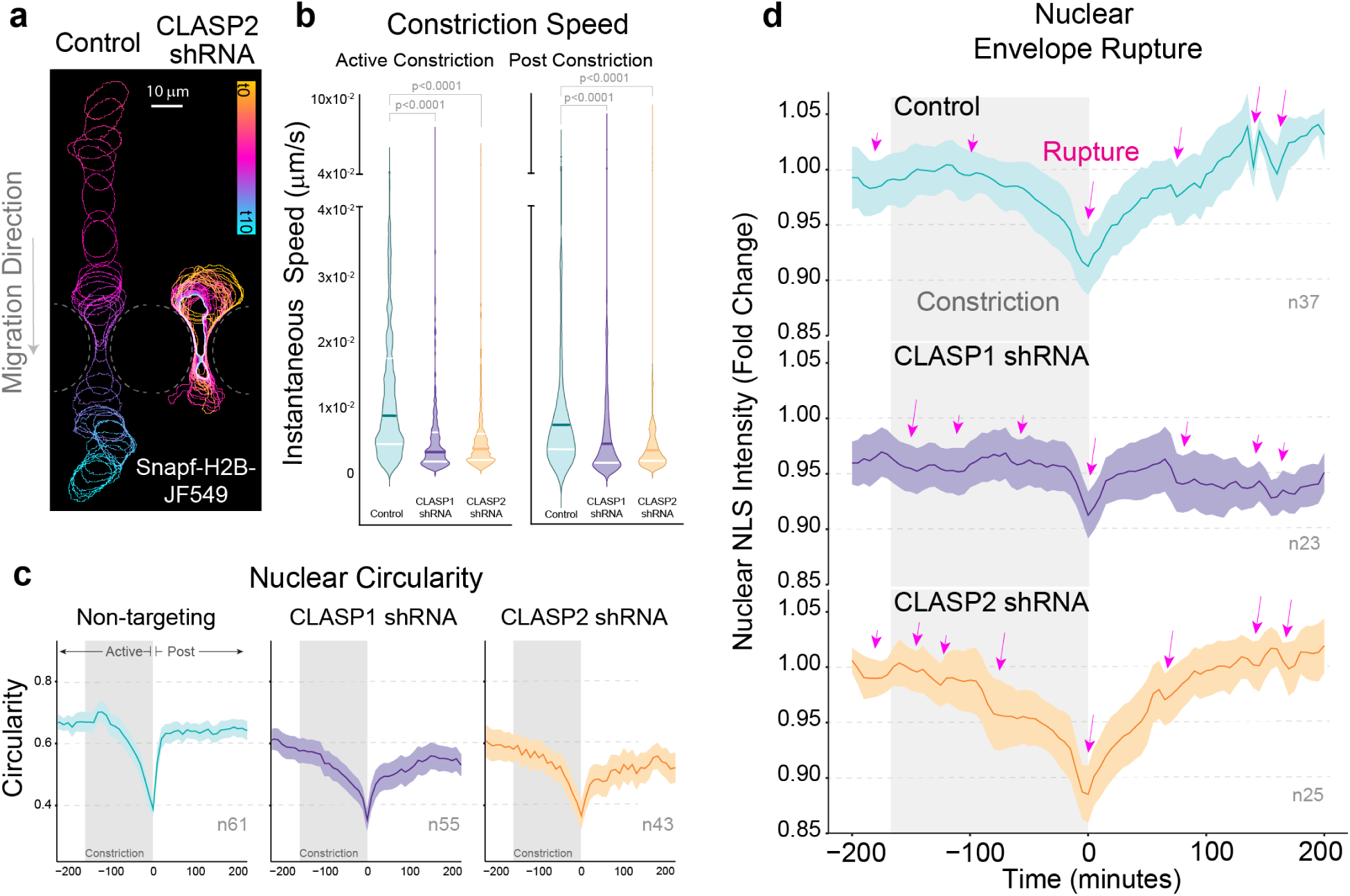
CLASP-depleted cells exhibit aberrant nuclear positioning and morphologies. (a) Temporal colour-coded projection of nuclear outlines of a control (non-targeting) and CLASP2-depleted cell undergoing nuclear constriction. (**b**) Instantaneous cell speeds during active and post nuclear constriction phases; comparing control (non-targeting) and CLASP2-depleted cells (CLASP2-shRNA). (**c**) Quantitation of nuclear circularity traces of control (non-targeting) and CLASP depleted cells showing lower nuclear circularity values as cells enter and exit constrictions. Graph shows mean circularity values with shaded 95% C.I. (**d**) Nuclear NLS intensity quantitation demonstrating major and minor rupture events (magenta arrows) during constriction. Graph shows mean with shaded 95% C.I.

**Extended Data Fig. 10.**
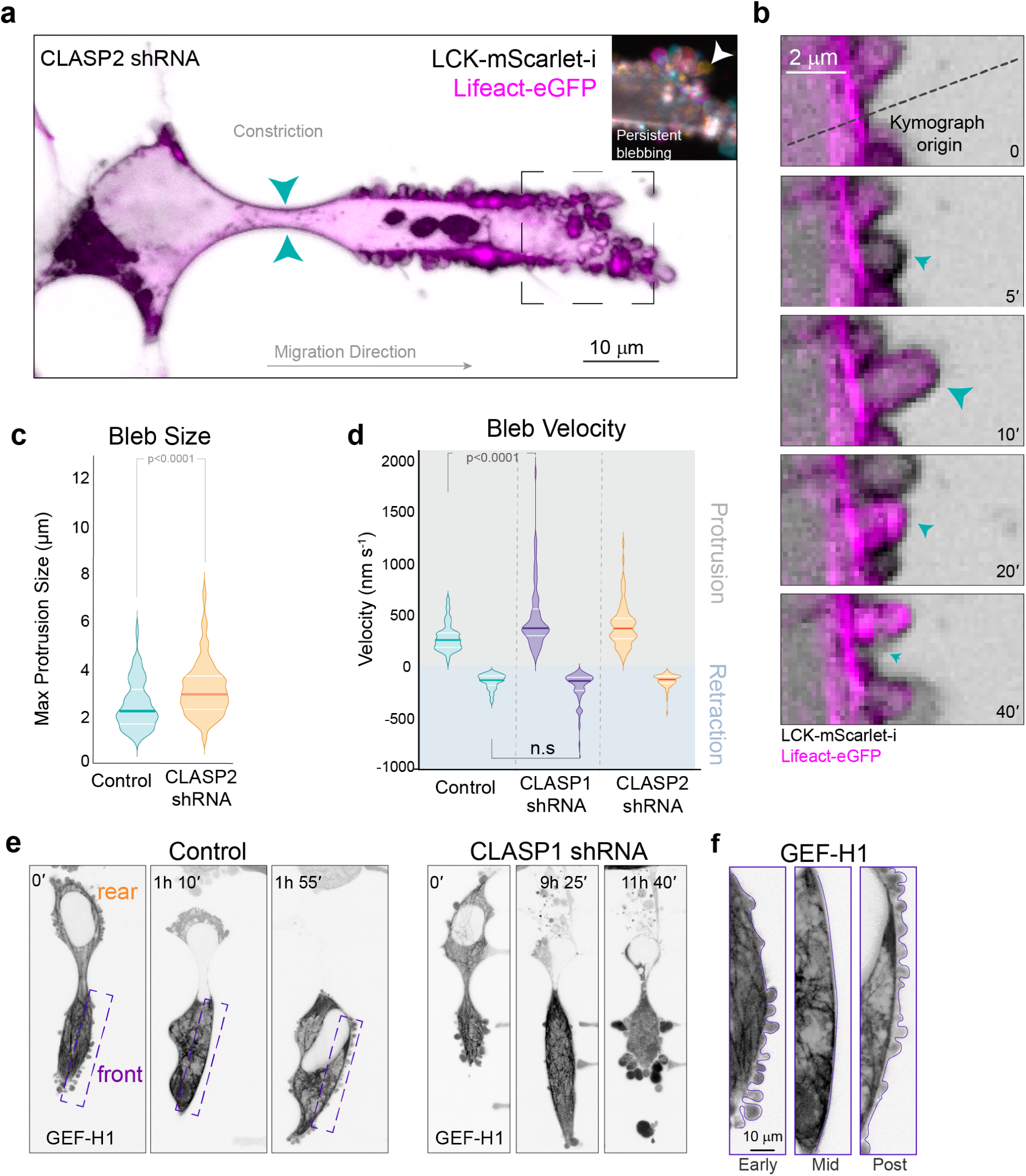
CLASP2-reinforced microtubules are required to coordinate contractility in confined cells. (**a**) Time-lapse images of CLASP-depleted (CLASP2-shRNA) cells co-expressing membrane (LCK-mScarlet-I) and cortical (eGFP-Lifeact) markers undergoing nuclear constriction in microchannels. Insets are temporal colour-coded projections (1 frame/sec for 1 min) of actin bleb dynamics at the cell surface. CLASP2-depletion increases bleb size (**b**) Enlarged inset of time-lapse imaging of bleb formation and recovery in a control cell. Membrane blebs (LCK, black) form from breaks in the actomyosin cortex (5 s) and are recovered by subsequent cortical repair (Lifeact-magenta; 10-20 seconds). Kymograph origin of Fig **4b**. (**c**) LASP2-depletion increases bleb size compared to control (non-targeting). (**d**) Analysis of bleb protrusion and retraction velocity. (**e**) GEF-H1 localization on microtubules in control (non-targeting) and CLASP1 depleted cells. (**f**) Enlarged insets from areas outlined in control cell front compartments in E (purple boxes) demonstrating changes in membrane blebbing in the front compartment during nuclear constriction phases.

**Extended Data Fig. 11.**
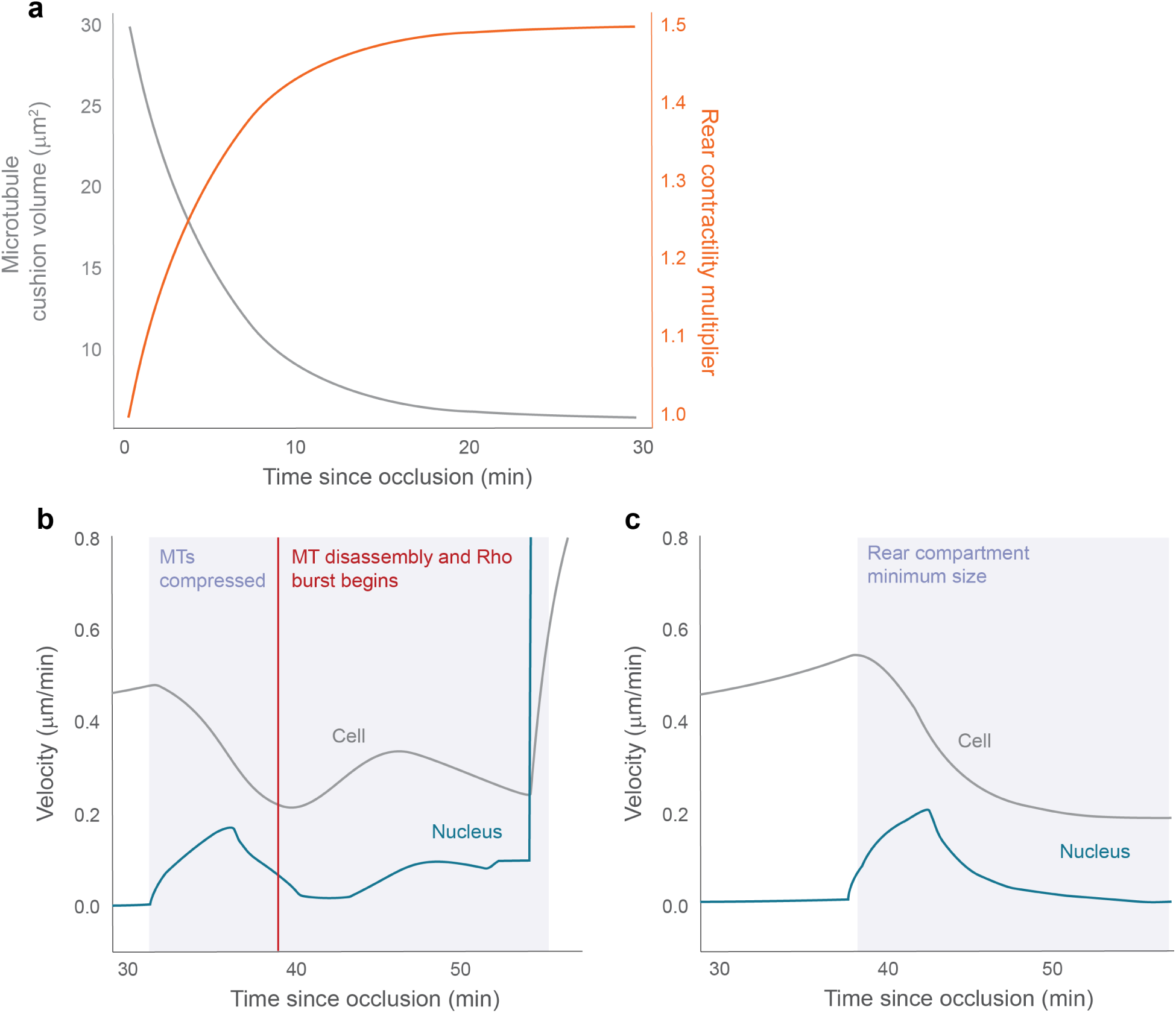
(**a**) Gradual shrinking of the rear microtubule cushion (grey line) and simultaneous increase of the Rho up-regulation factor (rear contractility multiplier (orange line) after initiation of microtubule cushion disassembly. (**b**) Velocity of cell (grey) and nucleus (blue) for the simulation in Fig. 6c and movie S10 depicting a control cell. The initiation of the Rho burst (red line) accelerates both cell and nucleus which finally leads to the sudden passage of the nucleus through the constriction. (**c**) Velocity of cell (grey) and nucleus (blue) for the simulation without the microtubule cushion in Fig. **6d** and Supplementary Video 11.

## Captions for Supplementary Videos

**Supplementary Video 1: Microtubules form a dynamic cage in cells moving through complex 3D environments.** Time-lapse Light Sheet Fluorescence Microscopy (LSFM) of 1205Lu CRISPR edited cells expressing meGFP-α-tubulin (orange) embedded into 2.5 mg/mL collagen I hydrogels within a custom-made sample holder. Collagen I was labelled and visualised using CNA35- mScarlet-I (cyan). Microtubules surround the nucleus in a single migrating cell, dynamically reorganising as the cell routes through collagen I fibres. Images were acquired every 30 seconds for a total duration of 2 hours, at each time interval 126 Z-slices were acquired at 0.4 µm steps totalling 50.4 um total. Images are Maximum Intensity Projections. Scale bar 10 µm.

**Supplementary Video 2: Contrasting microtubule organisation in 2D vs. 3D migrating cells.** Spinning disc confocal microscopy of 1205Lu CRISPR edited cells expressing meGFP-α-tubulin (orange) migrating on 2D glass coverslips and migrating within 2.5 mg/mL collagen I hydrogels. Collagen I was labelled and visualised using CAN35-Halotag-JF549 (cyan). Images of 2D cell migration was acquired every 1 minute for a total duration of 2 hours and 9 minutes. Images of 3D cell migration was acquired every 5 minutes for a total duration of 2 hours, at each time interval, 3 Z-slices were acquired at 3 µm spacing, totalling 9 µm total. Z-slices were Maximum Intensity Projected. Scale bar 10 µm.

**Supplementary Video 3: A microtubule cage assembles in cells undergoing constricted migration.** Rotating 3D-projection of 3D Structured Illumination Microscopy (3D-SIM) of cells undergoing constricted migration in microchannels, fixed and stained for acetylated (magenta), tyrosinated (grey) α-tubulin and nuclei (cyan). The microtubule cage structure can be seen to envelope the nucleus, lining the entire cell. Notably, in areas where the cell experiences active constriction, microtubules organise in parallel bundles surrounding the nucleus (mid). During early constriction timing (early) acetylated-α-tubulin is predominately localised to a posterior microtubule pool behind the nucleus (opposing the direction of migration) and interspaced between breaks of tyrosinated-α-tubulin. Z scroll through the rear compartment demonstrating density and curvature of microtubules posterior to the nucleus (24-26seconds). Scale bar 5 µm.

**Supplementary Video 4: CLASP depletion results in cell rupture during confined migration.** 1205Lu CRISPR edited cells co-expressing meGFP-α-tubulin (grey) and SNAPtag-H2B-JF549 (magenta). Control cells (left, non-targeting) successfully navigate constrictions between micropillars by deforming the nucleus and cell body. CLASP1 depleted cells (right) rupture. Images were acquired at 10-minute intervals for a total duration of 9 hours. Scale bar 10 µm

**Supplementary Video 5: Axial confinement results in microtubule depolymerisation in CLASP-depleted cells.** Live-spinning disc confocal time-lapse images of non-targeting control and CLASP1 depleted 1205Lu CRISPR edited cells expressing meGFP-α-tubulin undergoing compression (3 µm). Inverted to enhance contrast. Scale bar 10 µm.

**Supplementary Video 6: Axial confinement induces CLASP re-localisation**. Live-spinning disc confocal time-lapse images of 1205Lu cells expressing 2xmNeon-CLASP1 undergoing compression (3 µm) and release cycles. Upon release of compression, CLASP re- localises to +TIPs. Upon re-compression, CLASP binds along the lattice. Inverted to enhance contrast. Scale bar 5 µm.

**Supplementary Video 7: CLASP-depletion results in cortical blebbing.** Nested, time-lapse spinning disc microscopy of membrane (LCK-mScarlet-I, grey inverted) and F-actin (eGFP-LifeAct, magenta inverted) dynamics in non-targeting control and CLASP1 depleted 1205Lu cells migrating within constriction microchannels. Control cells exhibit blebbing at the rear cortico- membrane region during early- and mid-confinement stages. CLASP1-depletion evokes uncontrolled blebbing at both the anterior and posterior cortico-membrane regions. Every hour, for a total of 24 hours, nested acquisition of images were acquired at 1 second intervals for a total duration of 1 minute. Scale bar 5 µm.

**Supplementary Video 8: CLASP-depletion disrupts the proximal enrichment of myosin during confined migration.** Time-lapse spinning disc microscopy of non-targeting control and CLASP1 depleted 1205Lu CRISPR edited cells expressing eGFP-a-Tubulin (grey), and lentivirally transduced with mTurqoise2-MLC (myosin, cyan heatmap LUT) and SNAPtag-H2B- JF549 (magenta heatmap LUT). Control cells exhibit polarised proximal enrichment of myosin during early to mid-confinement stages. CLASP1-depletion perturbs polarised proximal myosin enrichment during confinement stages. Images were acquired at 15-minute intervals for a total duration of 24 hours. Scale bar 10 µm.

**Supplementary Video 9: Timed GEF-H1 activation upon microtubule depolymerisation precedes nuclear transmigration.** Time-lapse spinning disc microscopy showing ratiometric FRET/Donor images of non-targeting control and CLASP1 depleted 1205Lu cells lentivirally transduced with the GEF-H1-FLARE212 biosensor (Black-Purple-Yellow Heatmap LUT). Active GEF-H1 (high FRET/Donor ratio) can be visualised by yellow LUT colour and inactive (low FRET/Donor ratio) GEF-H1 in black-purple LUT colours. Active GEF-H1 polarises proximally in control cells during early to mid-constriction stages. CLASP1 depletion disrupts the proximal polarisation of active GEF-H1 causing cells to fail constricted migration and rupture. Images were acquired at 10-minute intervals for a total duration of 24 hours. Scale bar 10 µm.

**Supplementary Video 10: A timed Rho-burst at the cell-rear progresses nuclear transmigration during confined migration.** Time-lapse spinning disc microscopy of eGFP-RhoA (Black-Purple-Yellow Heatmap LUT) dynamics in non-targeting control and CLASP1 depleted 1205Lu cells. A polarised accumulation of proximal RhoA (yellow LUT colours) prior to nuclear translocation is associated with successful transmigration between constriction microchannels. CLASP1 depletion perturbs the polarised accumulation of RhoA, cells instead accumulate RhoA at both anteriorly and posteriorly leading to a static phenotype. Images were acquired at 10-minute intervals for a total duration of 24 hours. Scale bar 10 µm.

**Supplementary Video 11: Simulation of nuclear passage and the microtubule cushion.** The simulation of nuclear passage and simultaneous build-up of rear pressure (Fig. 6c) for a control cell (left). Simulation of failed nuclear passage (Fig. 6d) for a cell lacking the rear microtubule network (“cushion”) (right).

## Acknowledgments

We thank A.Yap, D.Bryant, T.Wittmann, S.Dumont, D. Barber, A.Molines and for discussion and comments on the manuscript. We also thank TRI Flow and Microscopy facility staff D.Khalil and Y.Ding, S.Roy and A.Ju, and IMB facility staff. We would like to acknowledge L.Lavis for his contribution of Janelia Fluorophores.

## Author contributions

S.J.S conceptualised and administered the project. R.J.J designed, performed and analysed all biological experiments with assistance from S.J.S, and M.D.W. A.D.F and D.B.O. performed mathematical simulations and developed the mathematical model with input from S.J.S R.J.J and M.D.W. S.J.S, R.J.J and M.D.W prepared data and schematics for visualization. P.T, M.N, K.M.D, G.D, and R.F. provided access to instrumentation, computing resources, analysis tools (lightsheet microscopes, FRET biosensors, data curation, software), and supervision. S.J.S, N.K.H. Funding acquisition. S.J.S and R.J.J wrote the original draft, S.J.S, M.D.W, A.J.L, K.M.D, G.D, N.K.H, and D.B.O. reviewed and edited the manuscript.

## Data and materials availability

All data are available in the main text or the supplementary materials. Plasmids are available on Addgene

