## Supplementary material for "Compression-dependent microtubule reinforcement comprises a mechanostat which enables cells to navigate confined environments": Mathematical Model

### Extended Data

### Full details of the mathematical model

#### Mechanical model for cell and nucleus

We introduce a mathematical model for single-cell transmigration between two micro-pillars. Our model describes the horizontal 1 $\mu$m thick cross-section of a cell migrating between two micro-pillars. The nucleus is treated as an elastic object represented by a closed contour line in the two-dimensional plane representing its nuclear envelope $\mathbf{n}(t,s) \in \mathbb{R}^2$. It is parametrised by $0 \leq s \leq 1$ and $t \geq 0$ represents time. Similarly, the cell cortex is represented by the closed contour line $\mathbf{m}(t,s) \in \mathbb{R}^2$. The entire setup is sketched in Figure 1.

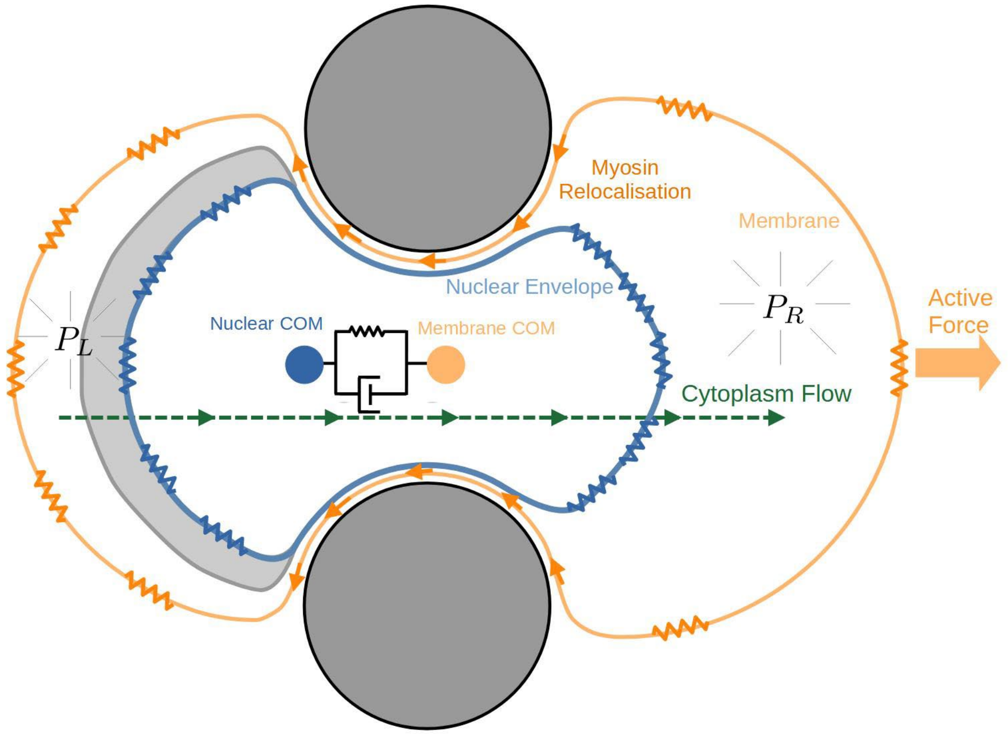


*Figure 1: Sketch of the mathematical model during occlusion: The active migratory force acts on both, the centers of mass of the membrane and – mediated by cytoskeleton fibers – of the nucleus. Surface friction is assumed to act on the cell's center of mass. It is coupled to the center of mass of the nucleus through a Kelvin-Voigt element modelling nucleus centering. The cell membrane and the nuclear envelope are under tension. The tension of the cortex adapts locally to the redistribution of myosin due to rearward actomyosin flow and diffusive exchange and pressurises the front and rear cytoplasm compartments. The resulting pressure differential between rear and front supports the transmigration of the nucleus and induces a flow of cytoplasm between the compartments which is stalled once the volume of the rear compartment reaches the volume associated with the microtubule cushion $V_{0,MT}^R$ shown in grey. Upon reaching a threshold pressure the MT cushion disassembles gradually to $V_{0,min}^R$). This triggers a simultaneous increase of Rho activity up-regulating rear tension which ultimately propels the nucleus through the constriction.*

We assume that the cell is exposed to a constant active migratory force in the direction of the passage between the two pillars (see Figure 2) written as $\mathbf{F}_{\text{migration}}$. The cell cortex and the nuclear envelope are both under tension which gives rise to the following stretching energies and associated forces,

$$\mathbf{F}^{\mathrm{C}}_{\mathrm{tension}} = \frac{\delta}{\delta \mathbf{m}} E^{\mathrm{C}}_{\mathrm{tension}} \quad \text{where} \quad E^{\mathrm{C}}_{\mathrm{tension}}[\mathbf{m}] = \sigma^{\mathrm{C}} \int_0^1 |\mathbf{m}'(s)| \; ds$$

and

$$\mathbf{F}^{\mathrm{N}}_{\mathrm{tension}} = \frac{\delta}{\delta \mathbf{n}} E^{\mathrm{N}}_{\mathrm{tension}} \quad \text{where} \quad E^{\mathrm{N}}_{\mathrm{tension}}[\mathbf{n}] = \sigma^{\mathrm{N}} \int_0^1 |\mathbf{n}'(s)| \; ds \; ,$$

where we model the stiffness of the nuclear envelope $\sigma^N$ and of the cell cortex $\sigma^C$ as constants and where $'$ represents the derivative with respect to $s$. When the nucleus obstructs the constriction between the two pillars we assume that the cortex is clamped on both sides between the nucleus and the pillars. In this case the rear and the front segment of the cortex are described by curves $\mathbf{m}^R = \mathbf{m}^R(t,s)$ and $\mathbf{m}^F = \mathbf{m}^F(t,s)$ and the stretching energy by

$$E^{\mathrm{M}}_{\mathrm{tension}}[\mathbf{m}^R, \mathbf{m}^F] = \sigma^R \int_0^1 |(\mathbf{m}^R)'(s)| \; ds + \sigma^F \int_0^1 |(\mathbf{m}^F)'(s)| \; ds \; ,$$

where $\sigma^R$ and $\sigma^F$ are the stiffness of the rear and front cortex.

The volumes (here: the area of the 2D cross-section) of the cytoplasm and of the nucleus are fixed and written as $V_0^C$ and $V_0^N$ respectively. When the nucleus occludes the passage the volume of the cytoplasm is divided into the volume of the rear compartment $V^R(t)$ and of the front compartment $V^F(t)$. In this case the stiffness of the cortex is assumed to vary between rear and front, written as $\sigma^R$ and $\sigma^F$, in response to the amount of myosin $M^R$ and $M^F$ in the cortex of the respective compartment. As a consequence, both compartments and also the nucleus are characterised by distinct pressure levels. They are defined by the constraints that the respective volumes are $V^R$, $V^F$ and $V_0^N$. The resulting forces enter the system of force balance equations, namely of forces acting on the cortex

$$\mathbf{0} = \mathbf{F}^{\mathrm{C}}_{\mathrm{pressure}} + \mathbf{F}^{\mathrm{C}}_{\mathrm{tension}} + \mathbf{F}^{\mathrm{C}}_{\mathrm{migration}} + \mathbf{F}^{\mathrm{C}}_{centering} + \mathbf{F}^{\mathrm{C}}_{\mathrm{friction}} + \mathbf{F}^{\mathrm{C}}_{\mathrm{steric}} \; ,$$

and of forces acting on the nuclear envelope,

$$\mathbf{0} = \mathbf{F}^{\mathrm{N}}_{\mathrm{pressure}} + \mathbf{F}^{\mathrm{N}}_{\mathrm{tension}} + \mathbf{F}^{\mathrm{N}}_{centering} + \mathbf{F}^{\mathrm{N}}_{\mathrm{steric}} + \mathbf{F}^{\mathrm{N}}_{\mathrm{migration}} \; .$$

This system of equations also involves forces due to nucleus centering which is modelled by a Kelvin-Voigt element (Hookean attractive force and drag in parallel) between the centers of mass of the nucleus and the cell, as well as repulsive forces (“steric”) preventing overlap between the nuclear envelope and plasma membrane/cortex and also between plasma membrane/cortex and micro-pillars. Note also that we assume that the migratory force $\mathbf{F}_{\mathrm{migration}}$ partially acts partially on the cortex ($\mathbf{F}^{\mathrm{C}}_{\mathrm{migration}}$) and on the nucleus ($\mathbf{F}^{\mathrm{N}}_{\mathrm{migration}}$).

**Actomyosin flux**

Once the nucleus occludes the passage between the micro-pillars we model two dynamic phenomena, myosin re-localization due to the rearward flow of actomyosin [29] and diffusive exchange between the compartments as well as the flow of cytoplasm past the nucleus.

The former is well-documented and plays a critical part in establishing a pressure gradient across the nucleus. To model it we introduce the fraction of cortical myosin contained in the rear and front compartments, $M^R = M^R(t)$ and $M^F = M^F(t)$, assuming that the total amount of cortical myosin is conserved, i.e., $M^R + M^F = 1$. We formulate a simple rate equation model stating that there is an influx of myosin into the rear compartment due to the rearward actomyosin flux which is proportional to the fraction of cortical myosin in the front compartment. We also include the effect of myosin exchange through the cytoplasm over or beneath the nucleus through an exchange term which is proportional to the difference of myosin fractions and the rate constant $\bar{\beta}$. This amounts to $\dot{M}^R = \beta M^F + \bar{\beta}(M^F - M^R)$ where $\beta > 0$ and $\bar{\beta} > 0$ are given. As a consequence – using that $M^R = 1 - M^F$–, the fraction $M^R$ follows the simple first order differential equation

$$\dot{M}^R = \beta(1 - M^R) + \bar{\beta}(1 - 2M^R) \ .$$

Finally, we assume that the stiffness of the cortex in the rear and at the front of the cell is given by a multiple of the fraction of cortical myosin,

$$\sigma^R = \gamma\mu M^R \quad \text{and} \quad \sigma^F = \gamma M^F,$$

where $\gamma > 0$ is a given constant and $\mu = \mu(t)$ is a dimensionless up-regulation factor which is initialised by $\mu_0 = 1$ and approaches $\mu_{\mathrm{Max}}$ (see below) upon initiation of rho up-regulation in the rear compartment.

Note that in order to keep the model simple we neglect the exchange of myosin between the two compartments as a consequence of the compartment boundary shifting while the nucleus is moving past the pillars.

### Cytoplasm flux

The total volumes of the nucleus and of the cytoplasm are assumed to be constant and given by $V_0^N$ and $V_0^C$. In situations when the nucleus occludes the passage between the micro-pillars dividing the cytoplasm into a front and rear compartment we explicitly distinguish between the volumes of the rear and front compartments, $V^R$ and $V^F$ in a way such that the total volume of the cytoplasm is conserved (Figure 2), $V^R + V^F = V_0^C$.

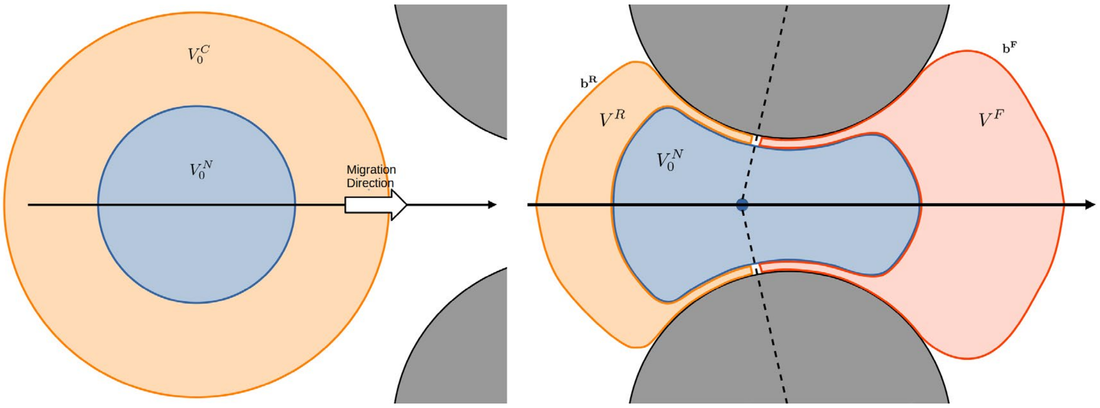


*Figure 2: We use a heuristic to identify occlusion of the passage between the two micro-pillars (grey) and to find the boundary points between front and rear (see below). This allows to split the volume of the cytoplasm $V_0^C$ into rear and front components $V^R$ and $V^F$.*

The redistribution of actomyosin triggers a difference in cortical tension and, subsequently, in intracellular pressure between the front and rear compartments. In agreement with the observations shown in Fig. 1 (main text), we assume that even after the nucleus occludes the space between the two pillars some cytoplasm still flows at a slow rate permeating the nucleus or else flowing beneath or above the nucleus. We model the flux of cytoplasm, typically from the rear into the front compartment, as proportional to the pressure difference between the two compartments,

$$\dot{V}^R = -\dot{V}^F = \alpha(P^R - P^F) ,$$

for a given constant $\alpha > 0$.

**Microtubule Cushion**

We also model a dense microtubule network in the rear compartment of the cell acting as a mechanical cushion with initial volume $V_{\mathrm{MT}}(t = 0) = V_{0,\mathrm{MT}}$, asserting a non-zero volume to this compartment.

This is designed to have no effect when the volume in the rear cytoplasmic compartment, $V^R$, is greater than the volume of the MT cushion, but will inhibit the volume of the rear compartment from shrinking below the volume of the MT cushion. We model this mechanical effect by stopping the decrease of $V^R$ when $V^R$ reaches the threshold value $V_{\mathrm{MT}}$.

We also model a mechanochemical effect assuming that once the pressure acting on the microtubule cushion (i.e. in the rear compartment) reaches the threshold level $P_{\mathrm{MT\ disassembly}}$ it gradually undergoes disassembly according to

$$\dot{V}_{\mathrm{MT}} = -\eta(V_{\mathrm{MT}} - V^R_{\mathrm{min}})$$

and simultaneously triggers an increase in Rho activity (Rho burst) up-regulating actomyosin contraction in the rear compartment of the cell. This is modeled as a gradual increase of the dimensionless up-regulation factor $\mu$ in the expressions for $\sigma^R$ and $\sigma^F$ according to

$$\dot{\mu} = \eta(\mu_{\mathrm{M}ax} - \mu) .$$

| **Symbol** | **Description** | **Value** | **Source** |
|---|---|---|---|
| $V_0^N$ | volume (area) of the nucleus | $50.3\ \mu\mathrm{m}^2$ | Heyden and Ortiz 2017 [30] |
| $V_0^C$ | volume (area) of the cytoplasm | $706.9\ \mu\mathrm{m}^2$ | Heyden and Ortiz 2017 [30] |
| $V_{\mathrm{MT}}$ | volume (area) occupied by rear MT network | $30\ \mu\mathrm{m}^2$ | |
| $V^R_{\mathrm{min}}$ | minimal volume (area) occupied by rear cytoplasm compartment | $6\ \mu\mathrm{m}^2$ | |
| $\sigma^N$ | nuclear envelope stiffness | $5000\ \mathrm{pN}\,\mu\mathrm{m}^{-1}$ | Nambiar, McConnell, and Tyska 2009 [31] |
| $\eta^C$ | surface friction | $10000\ \mathrm{pN\ min}\,\mu\mathrm{m}^{-1}$ | Tlili et al. 2018 [32] |

| | | | |
|---|---|---|---|
| $\mathbf{F}_{\text{migration}}$ | active migratory force | $(5000{,}0)^T$ pN | Lomakin et al. 2020 [33] |
| $\alpha$ | cytoplasm flow rate during occlusion | $0.01\ \text{min}^{-1}$ | |
| $\beta$ | actomyosin re-localization rate | $0.1\ \text{min}^{-1}$ | Venturini et al. 2020 [29] |
| $\overline{\beta}$ | random exchange of myosin | $0.1\ \text{min}^{-1}$ | estimated |
| $\kappa^N$ | elastic force for nucleus centering | $0.1\ \text{pN}\ \mu\text{m}^{-1}$ | estimated |
| $\eta^N$ | drag coefficient for nucleus centering | $0.5\ \text{pN min}\ \mu\text{m}^{-1}$ | estimated |
| $\gamma$ | coefficient for cortical tension | $5000\ \text{pN}\ \mu\text{m}^{-1}$ | Winklbauer 2015[1] |
| $\mu_{\text{Max}}$ | maximal coefficient for actomyosin up-regulation through Rho protein | 1.5 | main text, Fig. 5F |
| $\eta$ | rate of gradual MT cushion disassembly | $0.2\ \text{min}^{-1}$ | main text, Fig. 4I |

This finalizes the formulation of the mathematical model for the dependent quantities (degrees of freedom) $\mathbf{m} = \mathbf{m}(t,s)$, $\mathbf{n} = \mathbf{n}(t,s)$, $M^R(t)$ which also determines $M^F(t) = 1 - M^R(t)$ and $V^R(t)$ which determines $V^F(t) = V_0^C - V^R(t)$.

**Area and center of mass of regions enclosed by contour lines**

In this section we detail how the area enclosed by a closed contour line and its center of mass can be computed as path integrals and integrated into our mathematical model which is solely based on describing the surrounding contours of cell and nucleus. We denote the area of the region $\Omega$ enclosed by the closed curve $\partial\Omega = \{\mathbf{g}(s) = (g_x(s), g_y(s)), 0 \le s \le 1\}$ parametrised in anti-clockwise direction by $A[\mathbf{g}]$ and use the divergence theorem to compute it according to

$$A[\mathbf{g}] = \int_\Omega 1\ dx\,dy = \frac{1}{2}\int_\Omega \nabla \cdot \begin{pmatrix} x \\ y \end{pmatrix} dx\,dy = \frac{1}{2}\int_{\partial\Omega} \begin{pmatrix} x \\ y \end{pmatrix} \cdot \hat{n}\, d\ell = \frac{1}{2}\int_0^1 \left(x\, g_y{}'(s) - y\, g_x{}'\right) ds\ ,$$

where $d\ell = |\mathbf{g}'(s)|\, ds$ is the arc-length element and $\hat{n} = (g_y{}'(s), -g_x{}'(s))/|\mathbf{g}'(s)|$ is the unit outward normal.

Furthermore, we denote the center of mass in the $x$ direction of an area enclosed by the closed curve $\mathbf{g}(s)$ by $\mathbf{\Gamma}[\mathbf{g}] = (\Gamma_x[\mathbf{g}], \Gamma_y[\mathbf{g}])$ and compute it according to

$$\Gamma_x[\mathbf{g}] = \frac{1}{A[\mathbf{g}]}\int_\Omega x\; dx\, dy = \frac{1}{A[\mathbf{g}]}\frac{1}{2}\int_\Omega \nabla\cdot\begin{pmatrix}x^2\\0\end{pmatrix} dx\, dy =$$
$$= \frac{1}{2}\frac{1}{A[\mathbf{g}]}\int_{\partial\Omega}\begin{pmatrix}x^2\\0\end{pmatrix}\cdot\hat{n}d\ell = \frac{1}{2}\frac{1}{A[\mathbf{g}]}\int_0^1 x^2\, g_y{}'(s)\, ds\ ,$$

and an analogous expression for $\Gamma_y[\mathbf{g}]$.

**Determining occlusion**

We treat the passage between the micro-pillars as occluded when the distances between the center of mass of the nucleus and the micro-pillars are shorter than a threshold value (see Figure 2).

When the passage is occluded the intersections of the circular surfaces of the pillars with the straight lines connecting the center of mass of the nucleus and the center points of the pillars are considered the boundaries between the rear and the front compartments.

We use these boundary points to split the plasma membrane/cortex located at $\mathbf{m}(s)$ and the nuclear envelope located at $\mathbf{n}(s)$ into rear and front parts. Then we glue the rear parts of $\mathbf{m}$ and $\mathbf{n}$ together to obtain a closed contour written as $\mathbf{b}^R$ line enclosing the rear cytoplasm compartment and an analogous contour line written as $\mathbf{b}^F$ enclosing the front compartment.

Finally, we find the areas of the rear and front compartments using evaluating $A[\mathbf{b}^R]$ and $A[\mathbf{b}^F]$. Note that when the passage is not occluded the volume of the cytplasm can be computed evaluating the difference $A[\mathbf{m}] - A[\mathbf{n}]$.

**Numerical scheme**

We simulate the model sketched above using a splitting scheme to compute a time-discrete approximation to the solution $\mathbf{m}^n(s)$, $\mathbf{n}^n(s)$, $M^{R,n}$, $V^{R,n}$ with timestep $\Delta t$ and where we use $n$ as a discrete index for time. Note that for the contour lines of membrane/cortex and nuclear envelope we use a spatially discrete approximation of the functions $\mathbf{m}^n(s)$, $\mathbf{n}^n(s)$ , but we omit this detail in the following presentation.

1. In every timestep we update the closed contours for membrane/cortex and nuclear envelope, $\mathbf{m}^n(s)$ and $\mathbf{n}^n(s)$ according to an implicit Euler scheme for the force balance equations.
2. Then we compute separate updates for the amounts of actomyosin $M^{R,n}$ and $M^{F,n}$, namely an Euler step of their evolution equations and
3. an Euler step for the volume (area) of the rear compartment $V_0^{R,n}$ and utilizing that the volume of the front compartment is given by $V_0^{F,n} = V_0^C - V_0^{R,n}$.

The numerical treatment of the force balance equations is peculiar in that we compute an implicit Euler step by minimizing an associated energy functional which itself depends on the solution at the previous point in time,

$$(\mathbf{m}^n, \mathbf{n}^n) = \mathrm{argmin}_{\mathbf{m},\mathbf{n}} E_{\mathrm{total}}(\mathbf{m}^{n-1}, \mathbf{n}^{n-1}, M^{R,n-1}, V_0^{R,n-1})\ .$$

Note that for a minimiser $(\mathbf{m}^n, \mathbf{n}^n)$ the variation of $E_{\mathrm{total}}$ vanishes. Indeed, this energy functional is formulated in such a way that its variations with respect to $\mathbf{m}$ and $\mathbf{n}$ correspond to implicit Euler schemes for the force balance equations. Details about the variational formulation of a system of over-damped equations of motions can be found as previously published[2].

The energy functional

$$E_{\text{total}} = E_{\text{nucleus}} + E_{\text{cell}} + E_{\text{nucleus entering}} + E_{\text{steric}} \ ,$$

contains several components which refer to potential energies (corresponding to conservative forces), pseudo-energies (corresponding to drag forces) and penalizing potentials (corresponding to constraints). They are given by

$$\begin{aligned}
E_{\text{nucleus}} &= \frac{1}{\varepsilon}(A[\mathbf{n}^n] - V_0^N)^2 + E^{\text{N}}_{\text{tension}}[\mathbf{n}] - \nu^N \mathbf{F}_{\text{migration}} \cdot \Gamma[\mathbf{n}] \ , \\
E_{\text{cell}} &= \frac{1}{\varepsilon}\Theta[\mathbf{m},\mathbf{n}] + E^{\text{C}}_{\text{tension}}[\mathbf{m}^R, \mathbf{m}^F] + \\
&\qquad + \frac{\eta^C}{2\Delta t}|\Gamma[\mathbf{m}] - \Gamma[\mathbf{m}^{n-1}]|^2 - \nu^C \mathbf{F}_{\text{migration}} \cdot \Gamma[\mathbf{m}] \ , \\
E_{\text{nucleus centering}} &= \frac{\kappa^N}{2}|\Gamma[\mathbf{n}] - \Gamma[\mathbf{m}]|^2 + \frac{\eta^N}{2\Delta t}(|\Gamma[\mathbf{n}] - \Gamma[\mathbf{m}]| - |\Gamma[\mathbf{n}^{n-1}] - \Gamma[\mathbf{m}^{n-1}]|)^2 \ , \\
E_{\text{steric}} &= \frac{1}{\varepsilon}\int_0^1 (R - |\mathbf{c} - \mathbf{m}(s)|)_+^2 \, ds + \frac{1}{\varepsilon}\int_0^1 (-d_{\mathbf{m}}(\mathbf{n}(s)))_+^2 \, ds \ .
\end{aligned}$$

Note that friction with coefficient $\eta^C$ is acting on the center of mass of the cell, whereas the migratory force $\mathbf{F}_{\text{migration}}$ acts partially on the center of mass of the cell and of the nucleus according to the coefficients $\nu^C = 0.9$ and $\nu^N = 0.1$. This is to avoid un-physical negative pressure in the front compartment.

Constraints were implemented using steep penalization potentials whose coefficient written as $1/\varepsilon$ can be chosen arbitrarily large (though taking into account numerical stability).

Specifically, $\Theta[\mathbf{m},\mathbf{n}]$ is a penalizing potential that enforces $V_0^C$ as the total volume occupied by cytoplasm and $V^{R,n}$ and $V^{F,n} = V_0^C - V^{R,n}$ as volumes of the rear and front compartments during occlusion,

$$\Theta[\mathbf{m},\mathbf{n}] = \begin{cases} \frac{1}{2}(A[\mathbf{b}^R] - V^R)^2 + \frac{1}{2}(A[\mathbf{b}^F] - V^F)^2 & \text{channel is occluded,} \\ \frac{1}{2}(A[\mathbf{m}] - A[\mathbf{n}] - V_0^C)^2 & \text{otherwise.} \end{cases}$$

Also note that the first component in $E_{\text{steric}}$ penalizes overlap of plasma membrane and pillars according to a square potential. Here $(\ldots)+$ refers to the positive part of the expression enclosed in brackets. The second component penalizes intersections of nuclear envelope and membrane by evaluating the signed minimal distance $d_{\mathbf{m}}(\mathbf{x})$ of $x$ towards the closed curve representing the membrane $\mathbf{m}(s)$ (positive if $\mathbf{x}$ is inside, negative if it is outside).

Finally, the square penalization terms for the volume constraints in $E_{\text{cell}}$ and $E_{\text{nucleus}}$ deserve special mention. They are not only used to enforce the volumes of the nucleus and the cytoplasm, but also to compute the pressure in the rear and in the front compartment when the passage is occluded. These govern the flow of cytoplasm between compartments. They are computed as the (negative) variation of the penalization term $\Theta[\mathbf{m},\mathbf{n}]$ with respect to the compartment volumes

$$P_R = \frac{1}{\varepsilon}(V^R - A[\mathbf{b}^R]) \quad \text{and} \quad P^F = \frac{1}{\varepsilon}(V^F - A[\mathbf{b}^F]) \ .$$

**Simulations**

In our simulations the two micro-pillars correspond to two circles with radius $R = 14\ \mu$m centered at $\mathbf{c}^1, \mathbf{c}^2$. For simplicity we assume that they are aligned vertically and symmetrically with respect to the x-axis, at a distance of 2.5 $\mu$m. We also assume that the migratory force is horizontal, $\mathbf{F}_{\text{migration}} = (f_{\text{migration}}, 0)$, and that the nuclear envelope and the cell membrane are initially circular sharing a centre point on the horizontal axis. As a consequence, in simulations the cell approaches the micro-pillars horizontally. Indeed, with these choices of parameters and initial conditions, the simulation is symmetric with respect to the $x$ axis of the coordinate system. Exploiting this symmetry in simulations allows us to reduce the degrees of freedom by half.


**Acknowledgements**

Julia[3] was used to implement the numerical scheme and the Optim.jl package was used for its optimization part.
